# Microtubule Acetylation Regulates the Malignant Phenotype of Glioblastoma and is a Promising Therapeutic Target

**DOI:** 10.64898/2026.06.15.732342

**Authors:** Natanael Zarco, Athanassios Dovas, Santanu Bhattacharya, Tanmay Kulkarni, Misha Amini, Yi Liu, Ashley Haddock, Naveen KH Nagaiah, Debrabrata Mukhopadhyay, Paola Suarez-Meade, Alfredo Quinones-Hinojosa, Courtney A. Miller, Theodore M. Kamenecka, Peter Canoll, Rajappa S. Kenchappa, Steven S. Rosenfeld

## Abstract

Glioblastoma (GBM) is universally lethal despite decades of research to find effective treatments. This highlights the need to identify druggable targets essential for sustaining the malignant phenotype but dispensable for normal tissue. We propose that the enzyme α-tubulin acetyl transferase (*ATAT1*) meets these criteria. ATAT1 acetylates α-tubulin at lysine 40, which increases microtubule stability and promotes microtubule-based transport. While ATAT1 knockout mice have only a very mild phenotype, ATAT1 suppression in GBM has multiple therapeutic effects by reducing tumor invasion, proliferation, and therapeutic resistance. These translate not only into improved survival with ATAT1 targeting by itself, but also into synergy when ATAT1 deletion is combined with FDA approved therapies. This study strongly supports our conclusion that ATAT1 is a promising therapeutic target in GBM.

## INTRODUCTION

Microtubules (*MTs*) serve multiple roles in cell biology, including as tracks for MT transport motors; as regulators of cell mechanics and motility; and as central elements of the mitotic spindle and primary cilium^1–4^. These functions are also important for the malignant tumor cells^5–7^. This has motivated development of drugs that target MT polymerization and they have been very effective against a variety of malignancies^8,9^. However, given the importance of MTs in neuronal function and in mitosis, it is not surprising that these inhibitors also have significant neurologic and hematologic toxicities. An alternative would be to suppress MT function in a less toxic way.

One such approach would be to take advantage of MT post-translational modifications (*PTMs*). Referred to as the “tubulin code”, they include polyglutamylation, polyglycylation, tyrosination, detyrosination, phosphorylation, polyamination, and acetylation. These PTMs affect MT polymerization and its interactions with a variety of MT-binding proteins^10,11^. One of these PTMs is acetylation of K40 in the α tubulin subunit of the αβ tubulin heterodimer, the only PTM that occurs within the lumen of the microtubule. In vertebrates, K40 acetylation is catalyzed by one enzyme, α tubulin acetyl transferase (*ATAT1*)^12^. Acetylation of K40 increases MT flexibility, which prevents the MT lattice from fracturing and depolymerizing from the shear forces it experiences during normal cell function^13,14^. K40 acetylation also enhances intracellular transport by MT motors, cell motility, focal adhesion formation, and ligand activated receptor turnover^15–20^. Mice deleted for ATAT1 have no detectable α-tubulin acetylation in target tissues, indicating that other acetyl transferases cannot compensate for its loss^21^.

While the cytoskeleton, of which MTs are a central component, is essential for driving the malignant phenotype, it is also important for the function of normal cells. This has led to a widely held but untested assumption that targeting the cytoskeleton as an anti-cancer therapy would be unacceptably toxic. However, our recent work on the cytoskeletal motor non-muscle myosin II (*NMII*) has demonstrated that blocking NMII with a small molecule inhibitor is not only highly effective against GBM and but is also non-toxic^22^. Like NMII, ATAT1 drives many roles that are relevant to the malignant phenotype, suggesting that inhibiting it may have therapeutic benefits. Furthermore, ATAT1 knockout mice have only a mild phenotype which does not impair fertility or lifespan^21^, implying that an ATAT1 targeting strategy may have minimal toxicity.

In this report, we explore the contributions of ATAT1 to the malignant phenotype of GBM, among the most common and lethal of primary brain tumors. We find that ATAT1 is needed for GBM invasion, proliferation, oncogenic signaling, and therapy resistance. We also find that while depleting ATAT1 prolongs survival in murine models of GBM by itself, its targeting also synergizes with a variety of FDA-approved GBM therapeutics, including both MT depolymerizing drugs and temozolomide. These results demonstrate that ATAT1 is a promising therapeutic target in GBM.

## RESULTS

### ATAT1 regulates GBM cell morphology and mechanics

We shRNA suppressed ATAT1 in two human GBM cell lines (*L1 and 1A*), as well as in three murine GBM lines deleted for *Trp53*, *Pten*, or both (*referred to as Trp53*(-/-), *Pten*(-/-), or *Trp53/Pten*(-/-) *respectively*). The murine lines were developed using our genetically engineered murine models of GBM, in which a PDGF-IRES-cre retrovirus is orthotopically injected into the white matter of mice that have homozygous floxed alleles for *Trp53* and/or *Pten*. The resulting tumors are IDH wild type, PDGFRα driven, and deleted for *Trp53*, *Pten*, or both^23^. In both human and murine lines, ATAT1 KD reduces acetylated α tubulin by >85% (**Fig. S1A-C**). Compared to scrambled shRNA (*Scr*) transfected cells, ATAT1 knockdown (*KD*) cells are less polarized, as measured by an ∼2-fold increase in cellular circularity (**Fig. 1A,B**). We examined the effect of ATAT1 KD on the cellular content of polymerized MT by staining *Trp53/Pten*(-/-) GBM cells with SPY650-Tubulin, a conjugate of a fluorophore (SPY650^TM^) with docetaxel and counterstaining with Hoechst. At both high (**Fig. 1C**, *left*, *upper panels*) and low (**Fig. 1C**, *left*, *lower panels*) plating confluencies the normalized SPY650/Hoechst fluorescence ratio decreases by ∼50-60% with ATAT1 KD (**Fig. 1C**, *right*), with the most prominent changes seen in cortical MTs (**Fig. 1C**, *left*). MTs bind to and inactivate the Rho GTP exchange factor GEF H1, and MT depolymerization reverses this effect, leading to activation of RhoA and LIM kinase and consequent enhancement of actin polymerization and NMII contractility^24–26^. Consistent with this, ATAT1 KD increases stress fiber thickness and polymerized actin content by ∼50%, as measured by the normalized fluorescence of rhodamine phalloidin (**Fig. 1D & E**). Actin filament assembly and contractility in turn is a major determinant of cell stiffness^27–29^. This is consistent with our atomic force microscopy measurements (**Fig. 1F**), which show that ATAT1 KD increases Young’s modulus over the nucleus, the edge of the cell, and in between these two locations (*Nucleus, Periphery, and Midzone, respectively*) by 1.5-2-fold.

**Figure 1:**
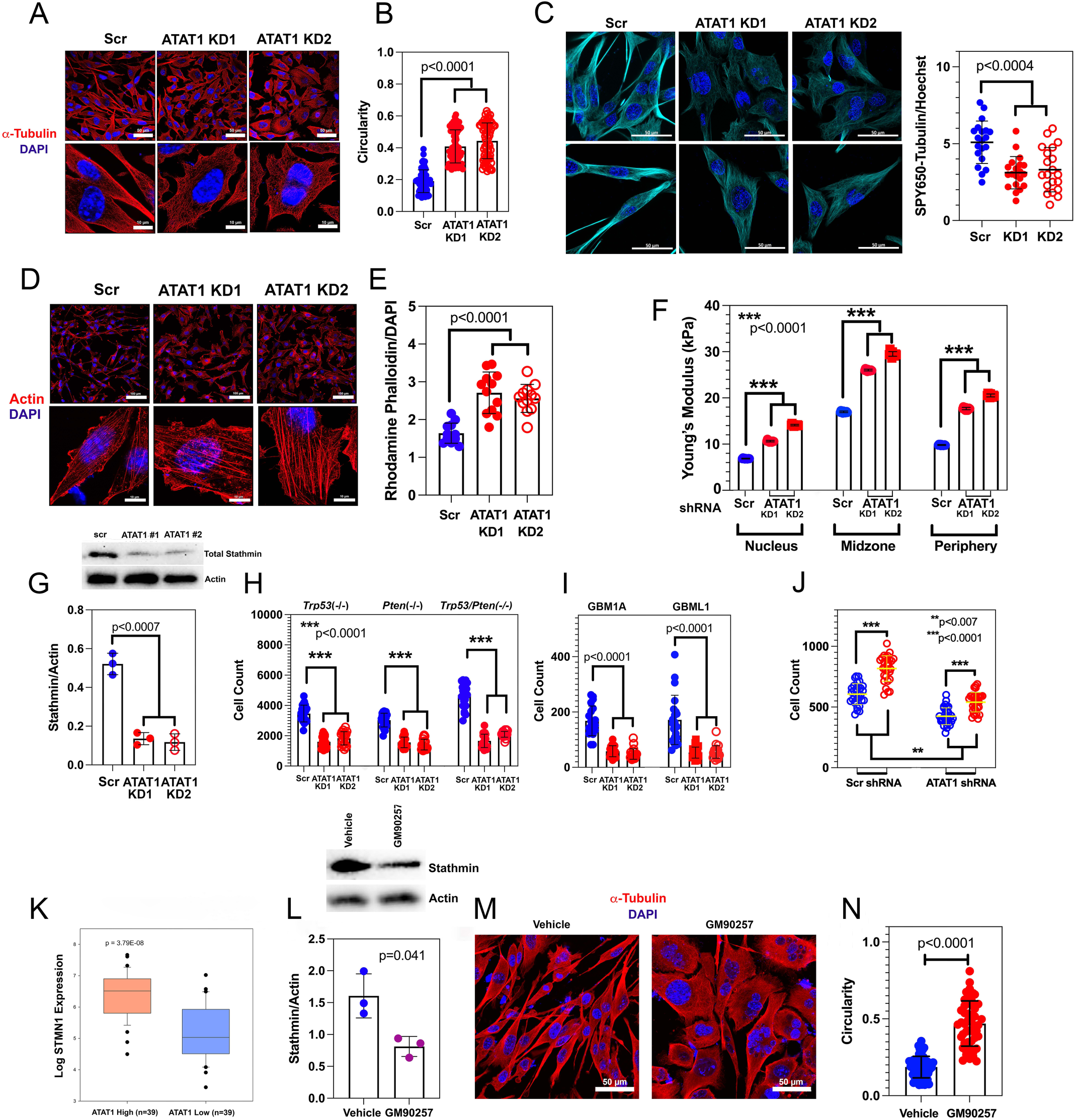
Tubulin acetylation regulates cell morphology and mechanics. (**A**): Staining of *Trp53/Pten*(-/-) cells for α tubulin (*red*) and DAPI (*blue*), transfected with either scrambled shRNA (*Scr*) or either of two ATAT1 shRNAs (*ATAT1 KD1, ATAT1 KD2*). Scale bars = 50 µm (*top*) and 10 µm (*bottom*). (**B**): Measurement of cell circularity for 60-80 cells from (**A**) demonstrates that ATAT suppression increases circularity ∼2-fold. (**C**): (*Left*) Staining of Scr, ATAT1 KD1, and ATAT1 KD2 treated *Trp53/Pten*(-/-) cells cultured at high (*upper*) and low (*lower*) confluency with SPY650-Tubulin and Hoechst. Scale bars = 50 µm. (*Right*) SPY650-Tubulin fluorescence normalized to Hoechst decreases ∼50-60% with ATAT1 KD. (**D**): Staining of *Trp53/Pten*(-/-) cells for actin with rhodamine phalloidin (*red*) and DAPI (*blue*), transfected with Scr, ATAT1 KD1, or ATAT1 KD2 shRNAs. Scale bars = 50 µm (*top*) and 10 µm (*bottom*). (**E**): Rhodamine phalloidin fluorescence normalized to Hoechst increases ∼50% with ATAT1 KD. (**F**): Young’s modulus of Scr and ATAT1 KD *Trp53/Pten*(-/-) cells measured over the perinuclear cytoplasm (*Nucleus*), cortex (*Periphery*), and in between (*Midzone*). (**G**): Knockdown of ATAT1 reduces total stathmin levels 3-4-fold. (**H,I**): Migration through 3 µm Transwell membranes is reduced 3-4-fold in murine *Trp53*(-/-), *Pten*(-/-), and *Trp53/Pten*(-/-) GBM lines (**H**) and in two human lines (GBM1A, GBML1) (**I**) with ATAT1 KD. (**J**): Transfection with a lentivirus encoding a GFP-stathmin fusion protein (*red*) increases *Trp53/Pten*(-/-) cell migration through a 3 µm Transwell membrane ∼50% compared a GFP encoding lentivirus control (*blue*) for both Scr and ATAT1-suppressing shRNA treated *Trp53/Pten*(-/-) cells. (**K**): Plot of data from TCGA demonstrates that stathmin mRNA expression is significantly higher in human GBMs with high ATAT1 expression than with low ATAT1 expression. (**L**): Treatment of murine *Trp53/Pten*(-/-) cells with 1000 nM GM902457 reduces stathmin expression ∼2-fold. (**M,N**): *Trp53/Pten*(-/-) cells were treated with vehicle or 1000 nM GM902457 and stained for α tubulin (*red*) and with DAPI (**M**). GM90257 treatment increases cell circularity ∼2-fold (**N**). See also **Figs. S1-S3**.

MT polymerization is regulated by stathmin, a 17 kDa protein that destabilizes MTs by sequestering αβ tubulin heterodimers. This allows cells to rapidly remodel their leading-edge during cell motility and invasion^30–32^. Stathmin is regulated by inactivating phosphorylation at several amino terminal serine residues, including S25. Knocking down ATAT1 leads to a 3-4-fold decrease in stathmin protein expression (**Fig. 1G**), with no significant change in the normalized content of pS25 stathmin (**Fig. S2**). These effects are associated with a ∼2-fold reduction in GBM cell invasiveness as measured by migration through 3 µm Transwell membranes in both murine (**Fig. 1H**) and human (**Fig. 1I**) lines. To further test the connection between ATAT1 KD, stathmin expression, and invasion, we suppressed ATAT1 in *Trp53/Pten*(-/-) cells and transfected these cells with either a GFP-encoding (*blue in* **Fig 1J**) or a GFP-stathmin-encoding (*red in* **Fig. 1J**) lentiviral plasmid. For both Scr and ATAT1 KD treated cells, transfection with a stathmin-encoding lentivirus increases Transwell migration ∼35%. To see if these findings carry over to the human setting, we queried the Cancer Genome Atlas (TCGA) for expression of stathmin mRNA (*gene name STMN1*) among GBM specimens representing the lowest and highest quartiles of ATAT1-expressing tumors. As **Fig. 1K** illustrates, the high ATAT1 group expresses nearly 3 times greater levels of STMN1 mRNA abundance than the low ATAT1 group (*p = 3.78 x 10^-^*^8^).

We also interfered with acetylation in a different way, by treating wild type *Trp53/Pten*(-/-) GBM cells with GM90257—a small molecule inhibitor that sterically blocks ATAT1 from accessing the K40 residue in α tubulin^33,34^. GM90257 inhibits acetylation with an EC_50_ of 274 ± 69 nM (**Fig. S3A**) and at a concentration of 1000 nM for 24 hours, it reduces stathmin levels ∼2-fold (**Fig. 1L**). Treatment with GM90257 increases cell circularity (**Fig. 1N** & **M**), similar to ATAT1 KD (**Fig. 1A** & **B**). We treated C57Bl6 mice with 25 mg/kg GM90257 given intraperitoneally every other day for 14 days and examined brain lysates for total and acetylated tubulin by Western blot (**Fig. S3B**). GM90257 has no appreciable effect on the brain content of K40 acetylated α tubulin compared to control, implying that it cannot cross the blood brain barrier (BBB) adequately to exert a pharmacologic effect.

### ATAT1 regulates sensitivity to MT depolymerizers

Vincristine and colchicine are MT depolymerizers that bind to distinct binding sites on the β tubulin subunit. Vincristine binds to the vinca site at the exposed “+” end, where it disrupts protofilament-protofilament interactions and induces MT disassembly; while colchicine binds to dissociated αβ tubulin heterodimers and keeps them from reattaching to the “+” end^35,36^. Thus, any condition that induces fracturing of the MT lattice, such as ATAT1 suppression, would be expected to generate new, dynamic MT ”+” ends. This predicts that variations in tubulin acetylation that occur naturally across different human GBM cell lines should correlate with sensitivity to any MT depolymerizers that bind to either the vinca or colchicine sites, and that suppression of ATAT1 should enhance the cytotoxicity of these drugs.

We measured the degree of tubulin acetylation in a set of primary human GBM cell lines and performed dose response studies on these cell lines with vincristine and BAL27862, the latter of which binds to the colchicine binding site. These human lines express levels of MT acetylation that span a range of >5-fold (**Fig. 2A & B**) and have EC_50_’s that span a range of ∼3 orders of magnitude (**Fig. S4A & B, Table S1**). The EC_50_ values for both vincristine and BAL27862 vary linearly with normalized acetylated α-tubulin content (**Fig. 2C & D**, *p=0.021, ρ=0.78 and p=0.011, ρ=0.82; respectively; Spearman’s rank correlation*). To test the second prediction, we suppressed ATAT1 in our murine *Trp53/Pten*(-/-) GBM line and measured the dose response to vincristine and BAL27862, along with two other MT depolymerizers, mebendazole, which binds to the colchicine site, and monomethyl auristatin E (MMAE), which binds to the vinca site. (**Fig. 2E-H**)^35^. In each case, ATAT1 KD shifts the dose response curves to the left >1.5 orders of magnitude (*p<0.0001, extra sum of squares F test; see also* **Table S2**), and we see similar findings using human 1A and L1 GBM lines (**Fig. S5 A-D** & **Table S4**). As a further control, we transfected ATAT1 intact, *Trp53*(-/-) murine GBM cells with an empty lentiviral plasmid(“*control plasmid*” in **Fig. 2I-K**), one encoding wild type ATAT1 (“*ATAT1 plasmid*” *in* **Fig. 2I-K**), or one encoding a catalytically inactive D157N ATAT1 mutant (“*mutant ATAT1*” *in* **Fig. 2K**). Transfection with either wild type or mutant plasmid increases immunodetectable ATAT1, but only the former enhances tubulin acetylation (**Fig. 2J**, *∼3-fold*). While transfection of wild type ATAT1 shifts the vincristine dose response curve to the right >100-fold (*p<0.0001, extra sum of squares F test*), transfection of the mutant D157N construct has no effect (**Fig. 2K**).

**Figure 2:**
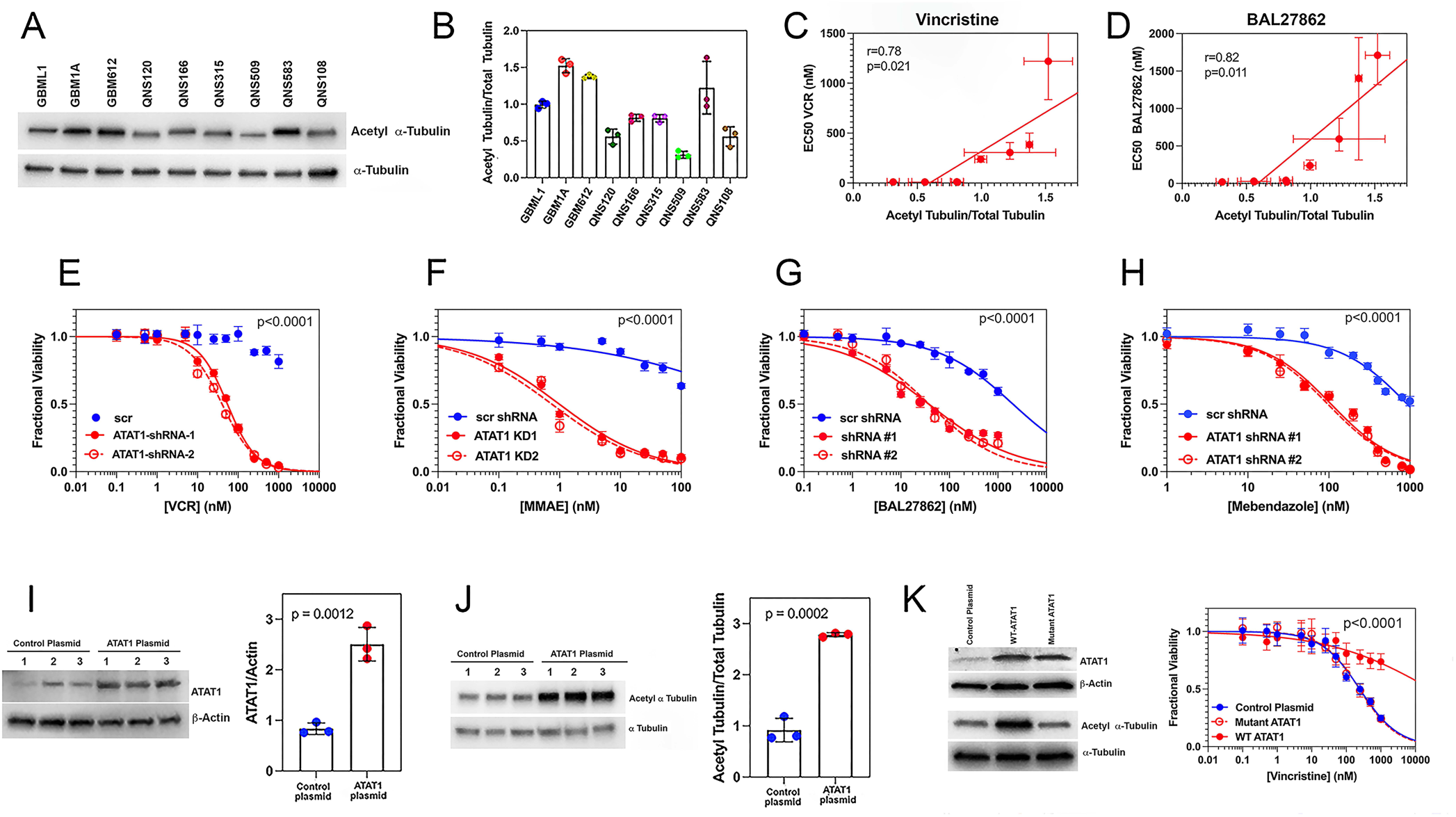
ATAT1 regulates sensitivity MT depolymerizers. See also Table S1. (**A** & **B**): Western blot (**A**) and quantitation of normalized expression of acetyl α tubulin (**B**) for 9 human GBM cell lines. (**C** & **D**): Plot of EC_50_, derived from dose response curves in **Fig. S4**, for human GBM cells treated with vincristine (**C**) or BAL27862 (**D**) *versus* normalized acetyl α tubulin content. Data were fit to a least squares linear regression with p values and goodness of fit indicated in the figure. Dashed grey curves represent 95% confidence intervals. (**E-H**): Dose response data of *Trp53/Pten*(-/-) cells treated with vincristine (**E**), MMAE (**F**), BAL27862 (**G**) and mebendazole (**H**) for cells transfected with Scr shRNA (*blue*) or ATAT1 encoding shRNAs (*red*). The corresponding EC_50_ values are derived from the dose response data in **Table S1** and are summarized in **Table S2**. Differences in values for ATAT1 KD and Scr EC_50_ are highly significant (*p<0.0001*). See also **Fig. S5** for corresponding dose response data for human 1A and L1 cell lines. (**I-K**): *Trp53*(-/-) cells were transfected with a control plasmid (**I,J**), one encoding ATAT1 (*ATAT1 plasmid*) (**I,J**), and one encoding a catalytically inactive D157N ATAT1 (*mutant ATAT1*) (**K**). While both ATAT1 and mutant ATAT1 plasmids increase expression of immunoreactive ATAT1, only the ATAT1 plasmid increases acetylated α tubulin. Transfection of *Trp53*(-/-) cells with the mutant ATAT1 plasmid (*open circles, red curve*) does not alter sensitivity to vincristine compared to the control plasmid (*blue*). By contrast, transfection with the ATAT1 plasmid (*solid red circles, red curve*) increases the EC_50_ of vincristine by more than two orders of magnitude (**K**). Differences in values for ATAT1 plasmid versus control plasmid transfected cells is highly significant (*p<0.0001*).

### ATAT1 regulates oncogenic signaling

MTs regulate signaling in several ways. First, they serve as tracks for dynein and kinesins, and acetylation significantly enhances the affinity and processivity of both motors^17,37–39^ and thereby increases trafficking of signaling molecules whose function requires active transport. This connection between trafficking and signaling is perhaps best illustrated by the primary cilium, a flexible “antenna-like” structure that functions as a signaling hub for pathways particularly important to malignant cells, including those downstream of Pdgfrα, Shh, Wnt, TGFβ, and Notch^40–44^. The primary cilium contains a central core of acetylated MTs, which serve as tracks for kinesin and dynein-mediated trafficking of signaling molecules into and out of the cilium^45^. Furthermore, by suppressing actin polymerization, MTs indirectly control the assembly and maturation of focal adhesions (FAs), whose signaling down the MAPK pathway can also be oncogenic.

To see how ATAT1 KD affects signaling in GBM, we performed bulk RNA-seq on Scr and ATAT1 shRNA treated murine GBM cells and applied DESeq to determine how ATAT1 KD affects gene set enrichment (**Table S3**). Differentially regulated gene ontologies for one of the two ATAT1 shRNA lentiviruses is depicted in **Fig. 3A**, and corresponding results using the other ATAT1 shRNA are shown in **Fig. S6**. The red dots in **Fig. S6** indicate gene ontologies shared by both datasets. Nearly all the differentially regulated ontologies shared in common are reduced with ATAT1 suppression, including those downstream of PI3 Kinase (*PI3K*), Hedgehog (*SHH*), STAT3, NFκB, NOTCH, and MYC. Enrichment plots for these pathways show that the downregulation of each of these is highly significant (**Fig. 3B**). A volcano plot (**Fig. 3C**, *blue, derived from* **Table S3**) demonstrates that ATAT1 KD downregulates key genes involved in proliferation and lineage determination, including Notch, Olig2, Sox2, CD44, Myc, and Ngfr. ATAT1 KD also downregulates genes that regulate inflammation, cell stress responses, and tumor invasion, including Fos, Jun, CCL2, and CD44.

**Figure 3:**
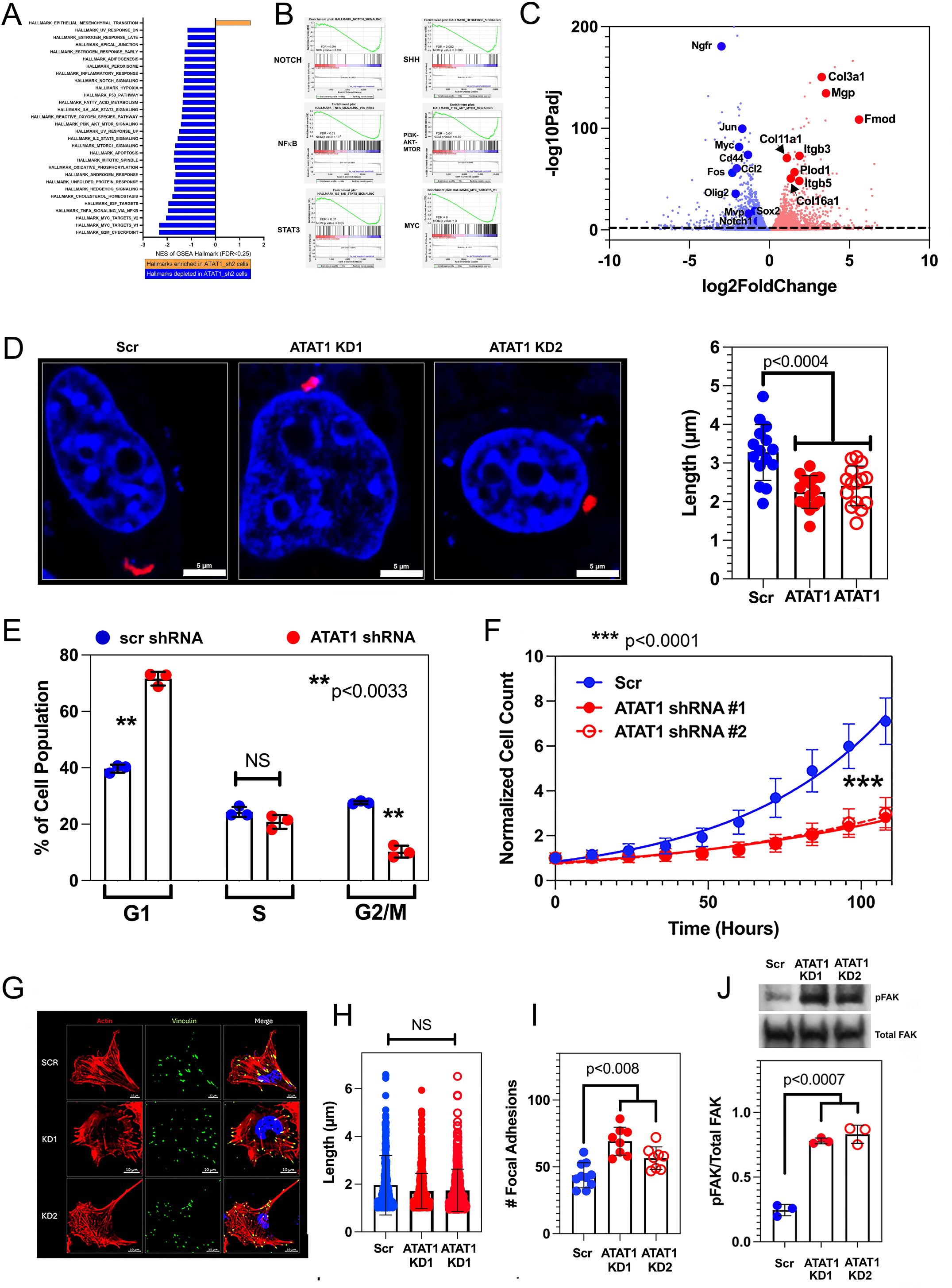
ATAT1 regulates oncogenic signaling. (**A** & **B**): GSEA (**A**) and enrichment plots (**B**) for gene sets differentially regulated with ATAT1 #1 shRNA knockdown. See **Fig. S6** for the corresponding GSEA plots for ATAT1 #2 shRNA. (**C**): A Volcano plot demonstrates downregulation (*blue*) of genes involved in supporting GBM proliferative capacity (NOTCH, OLIG2, SOX2, MYC, and NGFR) and stress/inflammatory response (FOS, JUN, CD44, CCL2); and upregulation (*red*) of genes involved in cell-ECM interactions (Itgb3, Itgb5) and ECM composition (Mgp, Fmod, Plod1, Col3a1, Col11a1, and Col16a1) (**D**): *Trp53/Pten*(-/-) cells were stained for the primary cilium marker ARL13B (*red*) and DAPI (*magenta*). ATAT1 KD shortens the primary cilium by ∼50%. (**E**): Flow cytometry of DAPI-stained Scr (*blue*) and ATAT1 (*red*) shRNA treated *Trp53/Pten*(-/-) cells shows that ATAT1 KD produces a partial block at the G_1_/S boundary. (**F**): Compared to Scr shRNA treatment (*blue*), ATAT1 (*red*) shRNA treatment of *Trp53/Pten*(-/-) cells increases doubling time 60-75%. Differences in growth rates between cells treated with Scr *versus* either of two ATAT1 suppressing shRNAs are highly significant (*p<0.0001*). (**G-I**): Murine *Trp53/Pten*(-/-) cells treated with Scr and ATAT1 KD shRNAs were stained for actin; vinculin, to visualize focal adhesions; and DAPI (**G**). While there is no significant change in mean length of focal adhesion length (**H**), the number of focal adhesions increases by ∼50% (**I**). (**J**): ATAT1 KD is associated with an ∼3-fold enhancement in the level of activated, pY397 FAK compared to Scr-treated cells. See also **Table S3**.

Given the role of the primary cilium in regulating oncogenic signaling, we examined how loss of ATAT1 affects ciliary structure by staining *Trp53/Pten*(-/-) cells for ARL13B. As **Fig. 3D** illustrates, suppression of ATAT1 reduces ciliary length ∼50% in *Trp53/Pten*(-/-) cells. Defective primary cilium function is frequently associated with corresponding defects in G_1_/S progression^47^, and we find that ATAT1 KD produces a partial G_1_/S block (**Fig. 3E**), associated with a >60% increase in doubling time (**Fig. 3F**; *34 ± 2 days for Scr shRNA treated, 56 ± 5 days and 61 ± 4 days for ATAT1 shRNA treated, p<0.0001*).

The Epithelial-Mesenchymal Transition ontology is the only one that is consistently upregulated with ATAT1 KD, and we identified the leading-edge genes that contribute to this enrichment (**Fig. 3C**, *red, derived from* **Table S3**). All of these are involved in interactions between the tumor cell and the extracellular matrix (*ECM*) and include those mediating cell ECM adhesion (*Itgb3, Itgb5*), composition (*Col3A1, Col11a1, Col16a1*), and modification (*Plod1, Fmod, Mgp*). Since focal adhesions (*FAs*) form an organizing hub for signaling downstream of cell-ECM interactions, we examined the effect of ATAT1 KD on FA length and number (**Fig. 3G-I**). While FA length is unaffected with ATAT1 KD (**Fig. 3H**), FA number increases by ∼50% (**Fig. 3I**). Focal adhesion kinase (*FAK*) connects ECM receptors, including integrins, to downstream signaling cascades, including the MAPK pathway, and ATAT1 KD is accompanied by an ∼3-fold increase in activated, pY397 FAK (**Fig. 3J**).

We validated the transcriptional effects of ATAT1 KD by first examining PDGFRα-dependent signaling—a key oncogenic pathway for GBMs with OPC and NPC lineages^48^. Transcriptional regulation of PDGFRA signaling in these subtypes can be regulated through NOTCH, FOS, JUN, MYC, OLIG2, and SOX2, which we have shown are all downregulated with ATAT1 knockdown (**Fig. 3A-C**). qRT-PCR shows that ATAT1 KD reduces PDGFRA mRNA ∼2-fold (**Fig. 4A**). There is also a highly significant correlation between ATAT1 and PDGFRα expression in human GBMs from the TCGA database (**Fig. 4B**). These transcriptional effects are mirrored in a ∼3-fold reduction in total and phosphorylated PDGFRα protein in both murine *Trp53/Pten*(-/-) cells (**Fig. 4C**) and in the human L1 and 1A GBM cell lines (**Fig. S7A**). To test if the reduction may be due to an acceleration of PDGFRα protein turnover with loss of α tubulin acetylation, we treated both control and ATAT1 KD cells with cycloheximide and monitored the disappearance of PDGFRα over time. Fitting the data to single exponential decays reveals that ATAT1 suppression *<u>slows</u>* PDGFRα turnover by ∼55% (**Fig. 4D**), consistent with the role of MTs in driving receptor internalization and trafficking^49–51^ and ruling out acceleration in PDGFRα turnover as an explanation. These changes are accompanied by a 2-3-fold reduction in activating phosphorylation of AKT and mTOR (**Fig. 4E** & **F**). As an additional control, we find that transfection of wild type murine GBM cells with a catalytically active ATAT1 plasmid nearly doubles expression of p-AKT (**Fig. 4G**). PDGFRα is not the only oncogenic receptor affected by ATAT1 KD, as loss of ATAT1 also significantly depletes NGFR at both the transcriptional (**Fig. 3C**) and protein levels (**Fig. 4H**). Pathways downstream of Notch, CD44, SHH, and STAT3 can stimulate expression of MYC, OLIG2, and SOX2^52–54^, and ATAT1 KD reduces protein and/or mRNA levels of all three of these transcription factors (**Fig. 4I-L**). MYC can bind to Aurora Kinase A (*AURKA*), which protects it from proteasomal degradation^55^, and a reduction in AURKA protein expression could therefore also explain our results. However, AURKA levels are unchanged by ATAT1 KD (**Fig. S7B**).

**Figure 4:**
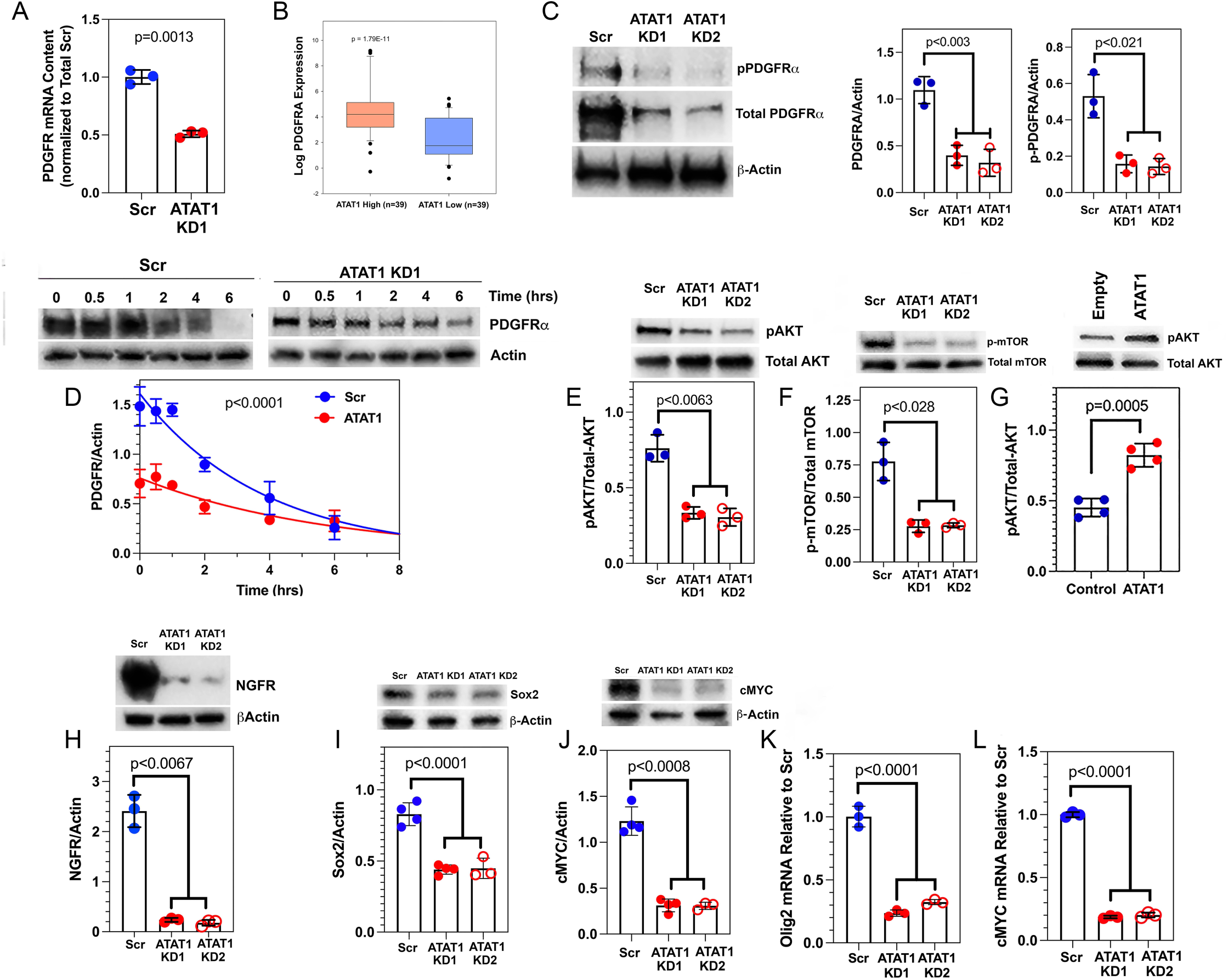
Knockdown of ATAT1 reduces oncogenic signaling along the PDGFRα signaling axis. (**A**): ATAT1 KD reduces PDGFRα mRNA, as measured by qRT PCR, by ∼2-fold. (**B**): Plot of data from TCGA demonstrates that PDGFRA mRNA expression is significantly higher in human GBMs with high ATAT1 expression than with low ATAT1 expression. (**C**): ATAT1 KD reduces expression of PDGFRα and pY849PDGFRα ∼3-fold in murine *Trp53/Pten*(-/-) murine GBM. Similar results were observed in human GBM1A and GBML1 cell lines as well (**Fig. S7**). (**D**): PDGFRα turnover was measured by blocking protein synthesis with cycloheximide and fitting the decrease in PDGFRα over time to single exponential decays. This reveals that ATAT1 suppression slows PDGFRα turnover by ∼55%. (**E** & **F**): ATAT1 KD reduces the content of activated pS473-AKT (**E**) and pT2446-mTOR (**F**) 2-3-fold. (**G**): Transfection of wild type murine GBM cells with a catalytically active ATAT1 expression plasmid nearly doubles expression of p-AKT. (**H**): ATAT1 KD reduces expression of NGFR >12-fold. (**I-L**): ATAT2 KD reduces protein levels of SOX2 (**I**) ∼2-fold anf of c-MYC (**J**) ∼4-fold and expression of OLIG2 mRNA (**K**) and c-MYC mRNA (**L**) 4-5-fold.

ATAT1 KD activates components of the MAPK pathway, including SRC, RAS, and ERK1/2 ∼2-3 fold (**Fig. 5A-C**), consistent with its effects on FAK (**Fig. 3J**). Transfecting wild type *Trp53/Pten*(-/-) cells with catalytically active ATAT1 has the opposite effect, reducing activated ERK1/2 ∼2-fold. (**Fig. 5D).** Tumor cells that upregulate oncogenic kinase pathways tend to become dependent upon them, a phenomenon known as oncogene addiction^56^, and can be killed with inhibitors of these pathways. We treated control and ATAT1 KD cells with inhibitors of SRC (*dasatinib*) and ERK1/2 (*ulixertinib*), and dose response data (**Fig. 5E** & **F**) show that ATAT1 KD enhances the potency of the former >100-fold and of the latter ∼10-fold (*p<0.0001 for both, extra sum of squares F test*). We also measured the levels of activated pT202/Y204 ERK1/2 and activated pY418 SRC in Scr and ATAT1 KD cells treated with either vehicle or the FAK inhibitor GSK2256098. FAK inhibition reduces levels of activated ERK1/2 to those of the scrambled control (**Fig. 5G**) and reduces corresponding levels of activated SRC by 75% (**Fig. 5H**).

**Figure 5:**
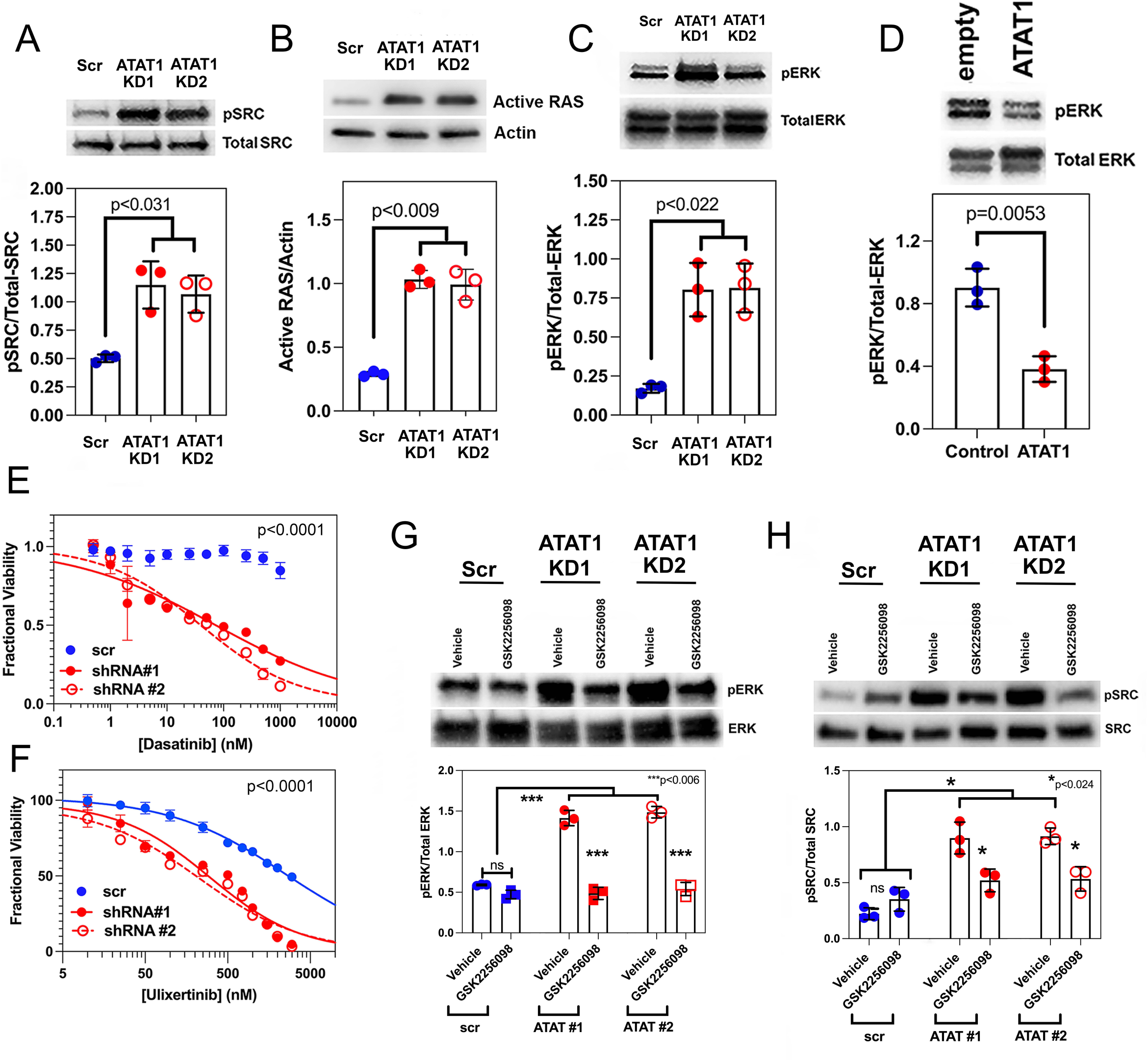
ATAT1 suppression activates the MAPK pathway through its effects on FAK. (**A-C**): ATAT1 KD increases expression of pY418 SRC (**A**), activated RAS (**B**), and pT202/Y204 ERK (**C**) 2-3-fold. (**D**): Transfection of wild type *Trp53/Pten*(-/-) cells with an ATAT1 expressing vector reduces ERK1/2 levels ∼2-fold. (**E** & **F**): ATAT1 KD is associated with a reduction in the EC_50_ for the SRC inhibitor dasatinib (**E**) and the ERK1/2 inhibitor ulixertinib (**F**). (**G** & **H**): The increase in pT202/Y204 ERK1/2 phosphorylation (**G**) and pY418 SRC phosphorylation (**H**) induced by ATAT1 suppression can be reversed by treatment with the FAK inhibitor GSK2256098.

### ATAT1 regulates sensitivity to temozolomide

Temozolomide (TMZ) is the standard alkylating agent for treating GBM, but its efficacy is limited by resistance. While the most important contributor to TMZ resistance is the dealkylating enzyme O⁶-methylguanine–DNA methyltransferase (MGMT)^58^, other mechanisms can support resistance as well. One of these is mediated by major vault protein (*MVP*). MVP is the main component of the vault ribonucleoprotein particle, a large, cytoplasmic, multi-subunit complex that sequesters anti-cancer therapeutics in the cytoplasm and enhances their export^59–61^. MVP transcriptional expression is regulated by FOS, JUN, and NF-κB, and mRNA expression of the former two and pathway activity downstream of the latter are reduced with ATAT1 KD, as is mRNA expression of MVP itself (**Fig. 3A-C**). Fos, Jun, and NFκB are also downstream of Notch and CD44^52,62,63^, and the latter two are downregulated with ATAT1 suppression (**Fig. 3A-C, Table S3**). These findings can explain why ATAT1 KD reduces MVP expression nearly 10-fold (**Fig.6A**). Signaling by Notch and CD44 depends on cleavage by γ secretase, and treatment of ATAT1 intact GBM cells with the γ secretase inhibitor LY411575 suppresses MVP expression by >2-fold (**Fig. 6B**).

**Figure 6:**
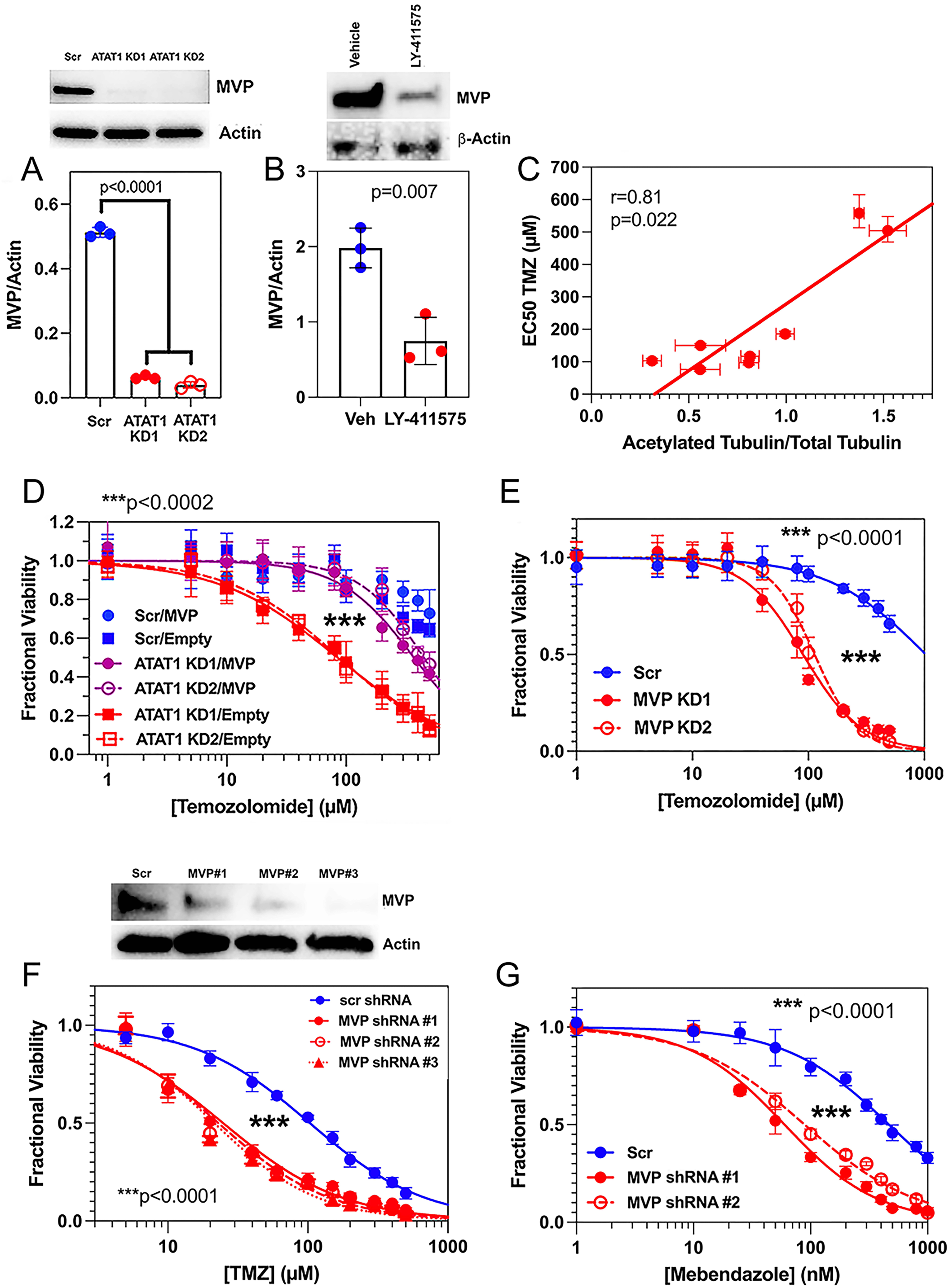
ATAT1 regulates sensitivity to temozolomide (see also Table S5). (**A**): ATAT1 KD reduces MVP levels >10-fold in *Trp53/Pten*(-/-) cells. (**B**): Treatment of *Trp53/Pten*(-/-) cells *in vitro* with 0.5 nM of the γ secretase inhibitor LY411575 for 24 hours reduces MVP expression >2-fold. (**C**). The EC_50_ for temozolomide varies directly with levels of normalized tubulin acetylation. (**D**): Dose responses to temozolomide for Scr or ATAT1 KD shRNA treated *Trp53/Pten*(-/-) cells that were transfected with a non-coding lentiviral vector (*empty*) or one encoding an EGFP-MVP fusion protein (*MVP*). (**E**): Suppression of MVP reduces temozolomide EC_50_ ∼13-fold in *Trp53/Pten*(-/-) cells. (**F**): shRNA suppression of MVP (*top panel*) reduces temozolomide EC_50_ >4-fold (*bottom panel*) in the temozolomide sensitive human QNS315 GBM cell line. (**G**): Suppression of MVP in *Trp53/Pten*(-/-) cells reduces the EC_50_ of mebendazole 5-7 fold.

If MVP provides some degree of TMZ resistance and if its expression depends on ATAT1 activity, we would predict that the EC_50_ for TMZ should vary with the degree of tubulin acetylation across a group of human GBM cell lines, akin to what we observed for vincristine and BAL27862 (**Fig. 2C,D**); and that it should increase or decrease with up- or down-regulation of ATAT1, respectively. We tested the first prediction by measuring TMZ dose responses for eight of the human cell lines described in **Fig. 2A** & **B**. Dose response curves are depicted in **Fig. S4C**, and a plot of TMZ EC_50_ *versus* degree of tubulin acetylation can be fit to a linear regression (**Fig. 6C**, *p=0.022, ρ = 0.81; Spearman’s rank correlation*). To test the second prediction, we upregulated MVP in ATAT1-KD GBM cells and downregulated it in wild type GBM cells and measured the effect on TMZ potency. ATAT1 or Scr shRNA-treated *Trp53/Pten*(-/-) murine GBM cells were transfected with either an empty lentiviral plasmid (“*Empty*” *in* **Fig. 6D**) or with one that encodes for an EGFP-MVP fusion protein (“*MVP*” *in* **Fig. 6D**). **Fig. S8A** documents the resulting increase in MVP expression. Scr shRNA-treated cells (*solid blue circles*) are highly resistant to TMZ (EC_50_∼1000µM, **Table S5**), and increasing MVP expression (*solid blue rectangles*) has little effect. While suppressing ATAT1 reduces the EC_50_ of TMZ >17fold (**Fig. 6D**, *red open and closed rectangles;* **Table S2**), resistance can be restored by forced expression of EGFP-MVP (**Fig. 6D**, *solid and open magenta circles*). Differences in the dose responses between the ATAT1 KD/Empty transfected cells (*red curves*) and ATAT1 KD/MVP transfected cells (*magenta curves*) are highly significant (*p<0.0002*, *extra sum of squares F test*). ATAT1 KD in the 1A and L1 human GBM cell lines also significantly enhances TMZ sensitivity (**Fig. S5E** & **F** *and* **Table S4**).

To test the importance of MVP to TMZ resistance in our GBM models, we suppressed its expression in *Trp53/Pten*(-/-) GBM cells by transfecting with either control shRNA or MVP-suppressing shRNA (**Fig. S8B**) and measured TMZ potency. As **Fig. 6E** and **Table S5** show, suppression of MVP reduces temozolomide EC_50_ ∼13-fold (*p<0.0001; extra sum of squares F test*), to a range that is generally considered to be TMZ sensitive, and which is similar to what we observe with ATAT1 suppression. To see if this effect on EC_50_ could also make a TMZ-sensitive tumor even more sensitive, we repeated this experiment on a human GBM cell line (QNS315) that at baseline is TMZ sensitive (EC_50_ 104 ± 5 µM, **Table S5**). MVP suppression in QNS315 lowers the EC_50_ by >4-fold (*p<0.0001*, *extra sum of squares F test*; **Fig. 6F, Table S5**).

Our proposal that tubulin acetylation regulates expression of PDGFR and MVP is based on the effects of ATAT1 KD and GM90257. To more rigorously test this proposed connection, we transfected Scr shRNA-treated and ATAT1 shRNA-treated *Trp53/Pten*(-/-) cells with either a non-coding plasmid or one encoding a K40 acetylation mimetic of α tubulin (K40Q). Previous studies have shown that expression of this pseudo acetylated α tubulin mimics the effect of tubulin acetylation on cellular contractility^83^ and reverses the effects of ATAT1 loss in *C. elegans*^84^. While transfection of Scr shRNA-treated control cells with the K40Q acetylation mimetic increases expression of PDGFRA and MVP by 10-15%, transfection of this construct in ATAT1 KD cells enhances expression of both 2-3-fold (**Fig. S9A-C**). As an additional orthogonal experiment, we transfected *Trp53/Pten*(-/-) cells with non-coding or K40Q α tubulin encoding plasmids and then treated these cells with vehicle or GM90257. Like ATAT1 KD, GM90257 suppresses expression of both PDGFRA and MVP, and this effect can be at least partially reversed with expression of the K40Q acetylation mimetic (**Fig. S9D-F**)

ATAT1 KD enhances the potency of mebendazole 9-fold (**Fig. 2H**, **Table S2**). While we have attributed this effect to MT fracturing and depolymerization with ATAT1 KD, reduction in MVP protein levels with ATAT1 KD might also enhance sensitivity to mebendazole, akin to what we see with TMZ. We measured the dose response of mebendazole in *Trp53/Pten*(-/-) cells treated with Scr or MVP suppressing shRNA. MVP KD increases mebendazole potency 5-8-fold (**Fig. 6G**, *p<0.0001, extra sum of squares F test***; Table S5**), which is very similar to the 9-fold increase in mebendazole potency that we observe with ATAT1 KD (**Fig. 2H**, **Table S2**).

### Targeting ATAT1 improves survival in pre-clinical GBM models

Our data show that ATAT1 has pleiotropic effects that in aggregate support GBM progression. These results predict that targeting ATAT1 should enhance survival in GBM and synergize with other therapies. To test this, we utilized two preclinical models. In the first, we shRNA suppressed ATAT1 in murine *Trp53/Pten*(-/-) GBM cells, orthotopically injected these cells into NSG recipients, and treated the mice with vehicle or with mebendazole. Both mebendazole alone (*25 mg/kg, 5 days/week by oral gavage,* **Fig. 7A**, *black versus blue*) and ATAT1 KD alone (**Fig. 7A**, *black versus magenta*) moderately increase median survival over the Scr shRNA, vehicle treated control (*17% and 31%, respectively; p<0.0002*). However, combining ATAT1 KD with mebendazole (**Fig. 7A**, *red*) increases median survival 91% over control, 61% over mebendazole alone, and 45% over ATAT KD alone (*p<0.0001 for all three comparisons*).

**Figure 7:**
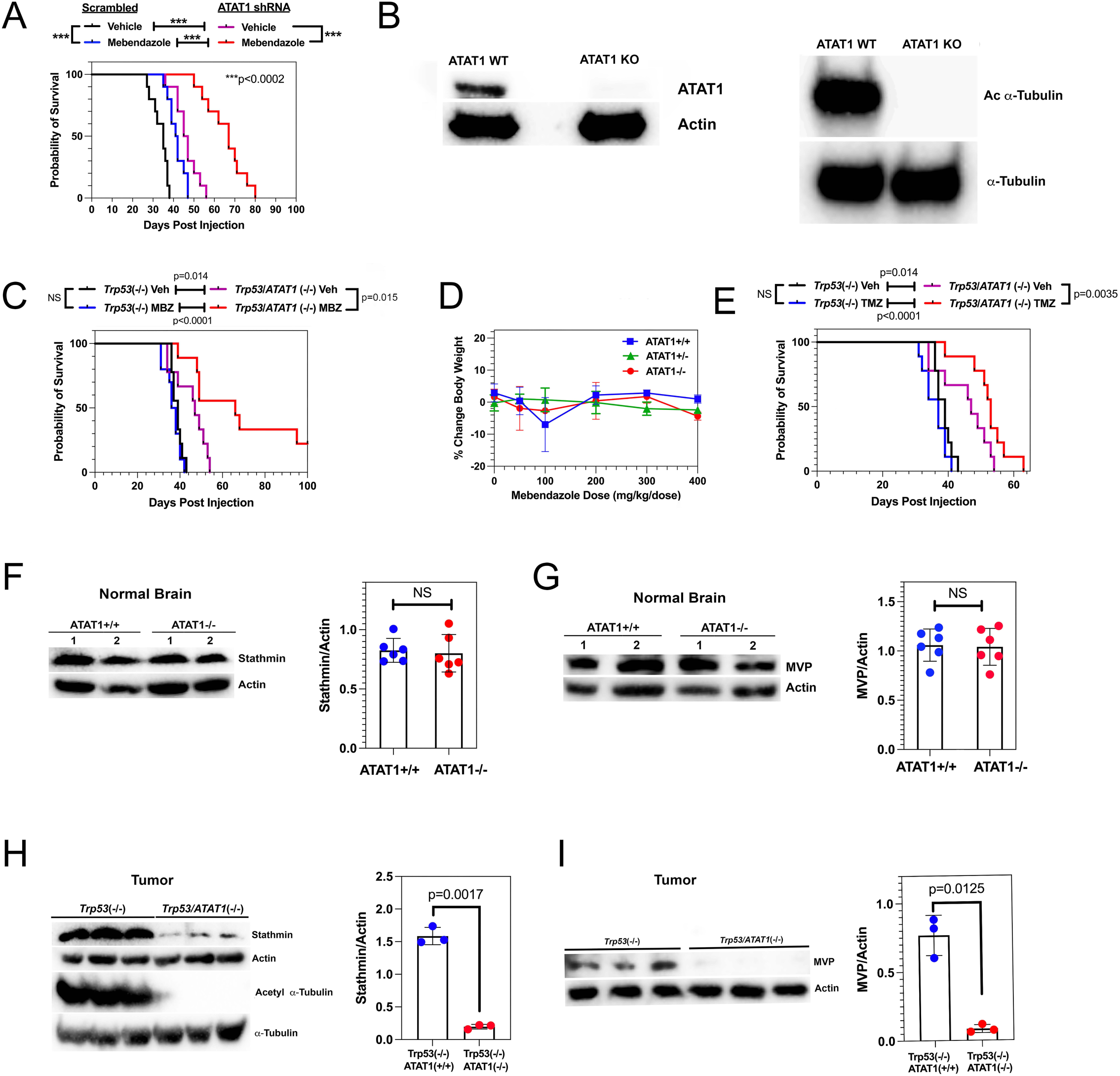
Targeting ATAT1 improves survival in pre-clinical GBM models. (**A**): Kaplan Meier survival curves for NSG mice orthotopically transplanted with murine *Pten/Trp53*(-/-) GBM cells treated with either Scr shRNA or ATAT1 shRNA. Compared to control, Scr shRNA treatment (*black*), ATAT1 KD (*blue*) moderately improves median survival for mice treated with vehicle. However, ATAT1 KD markedly improves median survival in the presence of mebendazole, compared to corresponding Scr shRNA control tumors. (**B**): Western blots for expression of ATAT1 (*left*) and acetylated α tubulin (*right*) in brain lysates from *Trp53* floxed mice that are intact (*ATAT1 WT*) or deleted (*ATAT1 KO*) for ATAT1. (**C**): ATAT1 deleted (*Trp53/ATAT1*(-/-)) and ATAT1 intact (*Trp53*(-/-)) GBMs were induced in the corresponding transgenic mice by orthotopic injection of a PDGF-IRES-cre retrovirus. Compared to mice with ATAT1 intact tumors treated with vehicle (Fig. 7C, *black*), mice with ATAT1 deleted tumors treated with vehicle (Fig. 7C, *magenta*) have median survival that is 21% longer (*p=0.014, log rank test*). While mebendazole treatment of mice with *Trp53*(-/-) GBM has no effect on survival compared to vehicle (Fig. 7C, *blue versus black*), the corresponding treatment of *Trp53/ATAT1* co-deleted GBM increases median survival by 40%, with 22% of mice surviving >100 days (Fig. 7C, *red versus magenta, p<0.0001*). (**D**). Single dose treatment of ATAT1 homozygous intact, ATAT1 heterozygous, and ATAT1 homozygous deleted mice with a single dose of 50, 100, 200, 300 or 400 mg/kg of mebendazole has no appreciable effect on weight at doses as high as 400 mg/kg. (**E**): *Trp53/ATAT1*(-/-) and *Trp53*(-/-) GBMs were induced in the corresponding transgenic mice by orthotopic injection of a PDGF-IRES-cre retrovirus and treated with temozolomide (TMZ) or vehicle. While TMZ treatment produces no survival benefit over vehicle in mice with ATAT1 intact tumors (*black versus blue*), it prolongs survival 13% compared to vehicle in mice with ATAT1 deleted tumors (*magenta versus red, p=0.04, log rank test*). Survival data for vehicle treated *Trp53*(-/-) and *Trp53/ATAT1*(-/-) GBMs are reproduced from Fig. 7C for comparison. (**F** & **G**): Western blots of stathmin (**F**) and MVP (**G**) from non-tumor bearing mouse brain show no appreciable effect of ATAT1 homozygous knockout on tissue levels of either. (**H** & **I**): The corresponding experiment on GBMs intact or homozygous deleted for ATAT1 show by contrast that in tumor, ATAT1 knockout markedly suppresses both stathmin (**H**) and MVP (**I**).

In our second model, we generated an immunocompetent transgenic mouse strain deleted for ATAT1 and containing homozygous floxed alleles for *Trp53*. Western blots demonstrate that brain lysates from these mice are devoid of both ATAT1 (**Fig. 7B**, *left*) and acetylated α tubulin (**Fig. 7B**, *right*), while brain lysates from *Trp53* floxed, ATAT1 intact mice contain both. We induced GBMs in these two genetically engineered mouse models (*referred to as Trp53(-/-) and Trp53/ATAT1(-/-)*) by orthotopic injection of our PDGF-IRES-cre retrovirus, began treatment with vehicle or mebendazole 5 days after tumor induction, and compared their survival. ATAT1 deletion by itself prolongs median survival 21% compared ATAT1 intact control tumors (**Fig. 7C**, *black versus magenta; p=0.014, log rank test*). While mebendazole treatment of mice with ATAT1-intact GBMs has no effect on survival compared to vehicle (**Fig. 7C**, *blue versus black*), treatment of ATAT1-deleted GBMs with mebendazole increases median survival by 40% over vehicle, with 22% of mice surviving >100 days (**Fig. 7C**, *red versus magenta, p<0.0001*). To determine if ATAT1 deletion increases mebendazole toxicity, we treated wild type, ATAT heterozygous deleted, and ATAT1 homozygous deleted mice (*designated ATAT1+/+, ATAT1+/-, and ATAT1-/-, respectively*) with a single dose of mebendazole and monitored weights on the day of dosing and every two days thereafter for two weeks. (*4 mice per group*). As **Fig. 7D** shows, mebendazole has no appreciable effect on weight after two weeks of treatment and at doses as high as 400 mg/kg. Our *in vitro* dose response studies show that *Trp53*(-/-) murine GBM cells are highly resistant to TMZ (**Fig. 6D** & **E, Table S5**). Consistent with this, mice with GBMs that are intact for ATAT1 experience no survival benefit from TMZ (**Fig. 7E**, *blue versus black*, *p=0.13, log rank test*). As noted above, ATAT1 deletion by itself increases median survival (**Fig. 7E**, *data for vehicle treated Trp53(-/-) and Trp53/ATAT1(-/-) GBMs reproduced from* **Fig. 7C**). By contrast, combining ATAT1 knockout with TMZ increases median survival of tumor bearing mice by 13% compared to vehicle treatment of ATAT1-deleted tumors (**Fig. 7E**, *magenta versus red*, *p=0.0035, log rank test*).

ATAT1 suppression of GBM cells *in vitro* markedly reduces levels of stathmin and MVP, respectively (**Figs. 1G** & **6A**). To see if this observation carries over into normal brain as well, we performed Western blots for stathmin and MVP on brains from ATAT1 intact (*ATAT1+/+*) and ATAT1 deleted (*ATAT1-/-*) mice. ATAT1 knockout has no significant effect on either stathmin or MVP expression in brain from ATAT1 knockout mice (**Fig. 7F** & **G**). We then repeated these experiments on orthotopic *Trp53*(-/-) and *Trp53/ATAT1*(-/-) GBMs that had been dissected from the brains of tumor bearing mice. By contrast, ATAT1 knockout in tumor markedly reduces expression of both stathmin and MVP (**Fig. 7H** & **I**).

## DISCUSSION

An effective therapeutic target in glioblastoma (GBM) must satisfy at least two criteria: it should drive multiple components of the malignant phenotype to minimize the ability of tumor cells to develop adaptive resistance, and it should be dispensable for normal tissue function to minimize toxicity. ATAT1 fulfills both requirements. MTs are necessary for many components of the malignant phenotype, including tumor invasion, oncogenic signaling, stress and inflammatory responses, and chemoresistance; and ATAT1 functions as an important modulator of how MTs regulate these programs. Furthermore, the proximal effects of ATAT1 KD on MT polymerization appear to induce additional responses that further enhance therapeutic potential. For example, while the downregulation of stathmin with ATAT1 KD mitigates against further MT depolymerization, it comes at the cost of impairing tumor cell invasion. Actin polymerization stimulates formation of focal adhesions^64^ while MT polymerization enhances FA turnover^82^, and ATAT1 KD increases the former and decreases the latter (**Fig. 1**). These reciprocal effects would be expected to stimulate focal adhesion assembly, which can impair cell motility^20^, and focal adhesion signaling. The former would decrease tumor cell invasion while the latter would increase MAPK signaling (**Fig. 5**). This latter effect would also lead to synthetic lethality when ATAT1 KD is combined with SRC or ERK inhibitors. Thus, ATAT1 loss not only suppresses some oncogenic pathways but also exposes new vulnerabilities through induced dependence on other pathways, a hallmark of oncogene addiction^56^.

ATAT1 also regulates therapeutic resistance. We have identified major vault protein (*MVP*) as a mediator of resistance that links ATAT1 to drug sensitivity for both TMZ and mebendazole. The MVP promoter contains FOS, JUN, and NFκB response elements^66^, and MVP transcription is upregulated under the inflammatory and cytotoxic conditions that rely on these transcription factors^67^. Notch activity increases MVP expression in breast carcinoma^68^, which may explain why downregulating Notch has been shown to enhance TMZ sensitivity^69^. The linear relationship between tubulin acetylation and EC_50_ for both MT depolymerizers and TMZ across a range of human GBM cell lines further supports a functional connection between cytoskeletal state and therapeutic sensitivity.

These *in vitro* effects of ATAT1 suppression translate into *in vivo* benefit. Knockdown or deletion of ATAT1 prolongs median survival in orthotopic murine GBM models by itself and synergizes with mebendazole. Mebendazole substantially prolongs GBM control in ATAT1-null mice, with a subset of animals achieving long-term survival, and it does so with no acute toxicity. Mebendazole is CNS-permeant and FDA approved, and it has been shown to prolong survival in pre-clinical models of GBM^70,71^. Furthermore, one of these studies^71^ showed that this drug is significantly more effective and less neurotoxic than vincristine; another MT depolymerizer that is part of a standard combination chemotherapy regimen for GBM^72^. ATAT1 KD also reduces the EC_50_ of temozolomide in our TMZ-resistant *Trp53/Pten*(-/-) murine GBM cell line *in vitro* to 80-90 µM. While this would explain why TMZ is significantly more effective in the setting of ATAT1 knockout, the magnitude of this improved survival is still relatively modest. One explanation for this difference is that the C_max_ for systemically administered TMZ in rodents is approximately 50-100 µM with corresponding values for brain 2-3-fold lower^73,74^. This means that for an ATAT1-deleted GBM model, peak TMZ concentrations would kill <30% of tumor cells, which would limit the robustness of a therapeutic response. However, MVP suppression also reduces the TMZ EC_50_ in the TMZ-sensitive QNS315 human GBM line (**Fig. 6F, Table S5**). This implies that targeting MVP has a meaningful biological impact regardless of the intrinsic sensitivity of a tumor to TMZ.

Stathmin and MVP expression remain unchanged in brain from ATAT1-null mice, whereas in ATAT1-null GBMs, expression of both is markedly reduced. This differential regulation underscores the point that the consequences of ATAT1 deletion depend on cellular context. Cytoskeletal regulators frequently exhibit cancer-specific dependencies due to chromosomal instability and synthetically lethal interactions. For example, chromosomally unstable tumor cells depend on the kinesin motor KIF18A for proliferation, while near-diploid cells do not^75^. Similarly, RB1-deficient cancer cells are hypersensitive to perturbations in microtubule stability^76^, and depletion of the microtubule regulator stathmin induces cell-cycle arrest and apoptosis in several cancer cell lines^77^. Such context dependence may explain why ATAT1 deletion is so well tolerated in mice while exerting profound effects in GBM.

Although the tubulin acetylation inhibitor GM90257 suppresses tumor growth in a flank xenograft model of triple negative breast cancer^33^, this compound has two significant limitations as a GBM therapeutic. First, it does not affect acetylated tubulin levels in normal brain, likely reflecting an inability to penetrate the blood brain barrier. Second, even if adequate CNS delivery could be achieved, the normal brain is rich in microtubules. Because the target of GM90257 is α tubulin and not ATAT1, a large fraction of this drug would be sequestered by the vast excess of microtubules in brain compared to tumor, buffering the availability of unbound drug for tumor cells. While direct ATAT1 inhibitors are not yet available, two reports provide a starting point for their development. First, a 1.05Å structure of the ATAT1 catalytic domain (*PDB: 4B5O*) provides a high-resolution map of the acetyl CoA binding site, the tubulin interacting interface, and the acetyl transferase catalytic residues^78^. Second, a more recent study used this structure to perform pharmacophore anchor modeling of ATAT1 and identified commercially available compounds, some FDA approved, with high calculated interaction energies, including ceftolozane, methotrexate, pemetrexed, cefonicid, and hesperidin^79^. These considerations suggest that the current lack of clinically optimized ATAT1 inhibitors should be viewed not as a barrier, but as an opportunity; and our findings provide a strong translational rationale for prioritizing development of brain-penetrant, selective ATAT1 inhibitors for the treatment of GBM.

## MATERIALS AND METHODS

### Kits

e-Myco TM Mycoplasma PCR Detection Kit (Lilif Diagnostics #25235), CellTiter-Glo (Promega #G9242), p24 ELISA kit (Clontech #632200), RNAqueous phenol-free total RNA isolation kit (Ambion, Life Technologies # AM1912), TURBO DNA-free™ kit (Ambion, Life Technologies # AM1907), Illumina TruSeq RNA Library Prep Kit v2 (Illumina), RNeasy Mini Kit (Qiagen # 74104), and iScript cDNA synthesis kit (BioRad #1708841). Active RAS pull down assay kit (Cell Signaling Technology # 11871/8821).

### Antibodies

anti–α-tubulin antibody (Cell Signaling Technology #2125), anti-Ki67 antibody (Abcam #ab15580), anti-MVP antibody (Protein Tech #16478-1-AP), anti-ARL13B antibody (Abcam #ab136648), anti-vinculin antibody (Sigma-Aldrich #V9131), anti-PDGFRa (Cell signaling technology #3164), anti-pY849 PDGFRa (Cell signaling technology #3170), anti-ERK (Cell signaling technology #9102), anti-pThr^202^/Tyr^204^-ERK (Cell signaling technology #4370), anti-mTOR (Cell signaling technology #2972), anti-pThr2446-mTOR (Cell signaling technology #63552), anti-AKT (Cell signaling technology #4691), anti-pSer473-AKT (Cell signaling technology #4060), anti-stathmin (Abcam #AB52630), anti-pSer25-stathmin, anti-ATAT1/C6ORF134 (Sigma Aldrich #SAB2108012), anti-FAK (Cell signaling technology #3285), anti-pY397-FAK (Cell signaling technology #3283), anti-SOX2 (Invitrogen #PA1-094), anti cMYC (Cell signaling technology #9402), anti-Aurora Kinase A (Invitrogen #PA5-97490), anti-NGFR (Abcam # 52987).

### Primers

Olig2 forward primer 5′-GGGAGGTCATGCCTTACGC-3′ (IDT Technologies), Olig2 reverse primer 5′-CTCCAGCGAGTTGGTGAGC-3′ (IDT Technologies), c-Myc forward primer 5′-AAAAAGCCACCGCCTACATC-3′ (IDT Technologies), c-Myc reverse primer 5′-ATGCACCAGAGTTTCGAAGC-3′ (IDT Technologies), and Prlp0 housekeeping gene primer (IDT Technologies). *Chemicals:* mebendazole (Selleck Chemicals #S4610), vincristine (Selleck Chemicals #S1241), temozolomide (Selleck Chemicals #S1237), Monomethyl auristatin E (MMAE) (MedKoo Biosciences #120202), Dasatinib (Selleck Chemicals #S1021), Ulixertinib (Selleck Chemicals #S7854), navitoclax (Selleck Chemicals #S1001), MCL1 inhibitor A1210477 (Selleck Chemicals #7790), puromycin (InvivoGen #ant-pr-1), sesame oil (Sigma Aldrich #S3547), DAPI (Thermo Fisher Scientific #62248), Hoechst 33342 (Thermo Fisher Scientific # H3570), 5% non-fat dry milk (BioRad #1706404).

### Reagents

DMEM+F12 media (Genesee #25-503), N2 supplement (Gibco #17502-048), hEGF (Sigma-Aldrich #E9644), hFGF (R&D systems #233-FB-025), NeuroPlex supplement (Gemini #400-161), Lipofectamine 3000 (Thermo Fisher Scientific #L3000008), DMEM media (Genesee #25-500), 10% FBS (LifeTechnologies #10437028), LentiX-Concentrator reagent (Takara Bio USA # 631231), rhodamine–phalloidin (Cytoskeleton Inc. #PHDR1), SPY650-Tubulin (Cytoskeleton Inc. #CY-SC503), and SYBR Green chemistry (Thermo Fisher Scientific #4367659). GM90257 was synthesized as previously described^33^. BAL27862 was graciously provided by Basilea Pharmaceutica.

### Plasmids

psPAX2 plasmid (Addgene #12260), pMD plasmid (Addgene #12259), pcDNA3-RFP (Addgene #13032), WT-ATAT1 pEF5B-FRT-GFP-αTAT1 (Addgene #27099), mutant-ATAT1 pEF5B-FRT-GFP-αTAT1 [D157N] (Addgene #27100), EGFP-MVP pEGFP-C2-MVP plasmid (Addgene #204543), stathmin-GFP (Addgene #86782), and pLV_TurboGFP-EV (Kenchappa, 2025). The lentiviral plasmid vector pLKO.1-puro based shRNA clones and control shRNA vector were purchased from Sigma-Aldrich: pLKO.1-puro non-targeting control SHC002, ATAT1 sh-RNA [TRCN0000193455 (ATAT1-shRNA-1), TRCN0000174781 (ATAT1-shRNA-2)] for mouse GBM cells, ATAT1 sh-RNA [TRCN0000167971 (ATAT1-shRNA-1), TRCN0000263600 (ATAT1-shRNA-2)] for human GBM cells, mouse MVP shRNA TRCN0000173596 (MVP-shRNA-1), mouse MVP shRNA TRCN0000175981 (MVP-shRNA-2).

### Mice

All mouse procedures were performed in compliance with the Mayo Clinic Institutional Animal Care and Use Committee guidelines (protocol numbers A00002923 and A00004179). Homozygous floxed *Trp53* mice in a C57Bl6 background (Stock #008462) were obtained from Jackson Laboratory. Studies were performed on mice between 7-12 weeks of age. The ATAT1 mouse strain used for this research project, C57BL/6N-*Atat1tm1(KOMP)Vlcg*/MbpMmucd, RRID:MMRRC_046712-UCD, was obtained from the Mutant Mouse Resource and Research Center (MMRRC) at University of California at Davis, an NIH-funded strain repository, and was donated to the MMRRC by The KOMP Repository, University of California, Davis; Originating from Kent Lloyd, UC Davis Mouse Biology Program. Mouse genotypes were regularly verified via tail snip (TransnetYX, Cordova, TN). Sample sizes were 10 mice per group in the Kaplan Meier survival experiments. Equal numbers of male and female mice were used in all these experiments, and mice were randomly assigned to each treatment group.

### Cell Lines

The human GBM line L1 was obtained from the laboratory of Dr. Justin D. Lathia (Lerner Research Institute of the Cleveland Clinic Foundation) and GBM lines 1A, 612, 120, 108, 166, 315, 509, and 583 were obtained from the BRIDGE Biobank of the Mayo Clinic Florida. The absence of *Mycoplasma* was confirmed regularly by e-Myco TM Mycoplasma PCR Detection Kit (Lilif Diagnostics, Cat# 25235) and cell lines were authenticated using STR analysis (IDEXX BioAnalytics).

### Glioma cell line isolation from mouse GBM tumor and culture

The protocol for isolation of tumor cells from *Trp53(-/-), Pten(-/-)* and *Trp53/Pten(-/-)* murine tumors has been described^23^. Human L1 primary GBM cell line was cultured and maintained in DMEM+F12 media with 1% N2 supplement (Gibco), 20ng/ml of hEGF (Sigma-Aldrich) and 20ng/ml of hFGF (R&D systems). Human 1A, 612, 120, 108, 166, 315, 509, and 583 primary lines were cultured and maintained in DMEM+F12 media with 1% NeuroPlex supplement (Gemini), 20ng/ml of EGF and 20ng/ml of FGF.

### Dose response curves/cell viability assays

5,000 cells/well were plated in 96-well plates and were allowed to attach for 48 hours. Cells were treated with various doses of mebendazole, vincristine, BAL27862, temozolomide, Monomethyl auristatin E (MMAE), Dasatinib, Ulixertinib, or vehicle for 72 hours and cell viability was measured using CellTiter-Glo (Promega, cat# G9242).

### Retrovirus production, intracerebral injections and drug treatment

PDGF-IRES-cre retrovirus was generated and injected intracranially according to methods described previously^23^. For the pharmacologic studies *Trp53^lox/lox^* or *Trp53*^lox/lox^/*ATAT1*(-/-) mice were injected intracranially with PDGF-IRES-Cre retrovirus, ten days later treated with vehicle, mebendazole (25 mg/kg by oral gavage, 5 days per week), or temozolomide (repeated cycle of 50mg/kg, daily for 5 days for 1 week with gap of 21 days), Treatment continued until tumor morbidity. NSG mice intracranially injected with *Trp53/Pten(-/-)* mouse GBM cells and treated with vehicle or mebendazole (25 mg/kg, 5 days/week by oral gavage) until tumor morbidity.

### Lentiviral production and transduction

Knockdown of ATAT1 in GBM cells was achieved via lentiviral infection with shRNA-encoding constructs. The lentiviral plasmid vector pLKO.1-puro based shRNA clones and control shRNA vector were purchased from Sigma-Aldrich (St Louis, MO, USA). The following constructs were used in these studies, non-targeting control (SHC002); ATAT1 sh-RNA [TRCN0000193455 (ATAT1-shRNA-1), TRCN0000174781 (ATAT1-shRNA-2)] for mouse GBM cells, ATAT1 sh-RNA [TRCN0000167971 (ATAT1-shRNA-1), TRCN0000263600 (ATAT1-shRNA-2)] for human GBM cells Each of the pLK0.1 targeting constructs were cotransfected with psPAX2 and pMD plasmids into HEK-293T cells via Lipofectamine 3000 transfection agent (Life Technologies, catalog # 11668027) in serum-free medium. After 8 hours of transfection, the viral particle-containing medium was removed and replaced with fresh complete medium. Transfected cells were then grown in DMEM media containing 10% FBS for 48 hours at 37°C, 5% CO2. Media containing virus was harvested and centrifuged for 10 mins in a clinical specimen centrifuge and then filtered through a 0.45 μm filter. Lentiviral particles were concentrated using LentiX-Concentrator reagent (Takara Bio USA) and the viral titer was determined using a p24 ELISA kit (Clontech). Human GBM cells were infected by incubating with virus containing media (10 MOI of virus and 4μg/mL of polybrene (Sigma-Aldrich)) overnight. Cells were selected for positive shRNA infection using puromycin (0.5ug/ml) for seven days and maintained in 0.1ug/mL puromycin containing media.

### Transfection of GBM cells

GBM cells were plated to 70% confluency in 6 well plates, chamber slides or 96 well plates. 24 hours after plating, cells were transfected using Lipofectamine 3000 transfection reagent (Invitrogen) according to the manufacturer’s instructions. Briefly, pcDNA3-RFP, WT-ATAT1, mutant-ATAT1, EGFP-MVP, stathmin-GFP, shRNA plasmids targeting mouse MVP gene were mixed with Lipofectamine 3000 reagent. Then p3000 reagent was added, mixed, and incubated at room temperature for 20 minutes and transfection mixture was added to the cell culture, which was incubated for 48-120 hours for biochemical assays.

### Transwell invasion assay

Fluoroblok Transwell inserts added with 100,000 GBM cells for each insert. 10% FBS was used as a chemoattractant. Cells were incubated for 12 hours at 37°C, inserts were washed with PBS, fixed in 4% PFA for 15 min and washed twice with PBS before staining with DAPI. Images were captured and analyzed for nuclear counts using Cytation 5.

### Western blot analysis

Cells were scraped and incubated in lysis buffer (50 mM Tris HCl at pH 7.4, 150 mM NaCl, 1 mM EDTA, 1.0% Nonidet P-40, and a mixture of protease and phosphatase inhibitors), on ice from 30 minutes. Debris was removed by centrifugation for 10 minutes at high speed at 4°C, and cleared lysates were run on SDS/PAGE and transferred to polyvinylidene difluoride membranes. Membranes were blocked in 5% non-fat dry milk in TBS + 0.1% Tween 20 for 1 hour at room temperature, incubated with primary antibody in blocking solution for overnight at 4°C, followed by secondary antibody for 1 hour at room temperature, and developed using an enhanced chemiluminescence solution.

### Mebendazole treatment of ATAT1 deleted mice

Wildtype, ATAT1+/- or ATAT-/- C57Bl/6J mice were single dosed orally with mebendazole at 50, 100, 200, 300 or 400 mg/kg or vehicle (sesame oil). Mice were weighed on the day of dosing and every two days for two weeks.

### Bulk RNA-seq data acquisition and analysis

RNA sequencing was performed at the Columbia Sulzberger Genome Center. Total RNA from three independent biological replicates (naïve and ispinesib resistant cells) was isolated using the RNAqueous phenol-free total RNA isolation kit (Ambion, Life Technologies, Grand Island, NY) and DNA contamination in isolated RNA was removed by DNase treatment using TURBO DNA-free™ kit (Ambion, Life Technologies, CA). All samples had an RNA Integrity Number greater than 7.6, as assessed using Agilent Bioanalyzer. Libraries were prepared using the Illumina TruSeq RNA Library Prep Kit v2 and 20 million paired-end, 75 bp reads were acquired on an Element Aviti sequencer at Columbia Genome Center. Reads were pseudoaligned to a kallisto mouse transcriptome index (GRCm38.p6) using kallisto (0.44.0), and differential gene expression analysis was determined using DESeq2. Gene Set Enrichment Analysis (GSEA) was performed on the desktop version of GSEA (v4.1.0), using Hallmarks and the Verhaak_Glioblastoma_Mesenchymal gene sets from the Molecular Signatures Database (MSigDB).

### Measurements of Young’s Modulus

AFM experiments were performed using the Dimension FastScan system equipped with the ScanAsyst™ mode and an ICON head (Bruker Corporation, Santa Barbara, CA). To investigate cellular nanomechanical properties, we utilized Peak Force Quantitative Nanomechanical Mapping Live Cell probe version 2.0 featuring spherical tip geometry. The probe’s nominal spring constant and tip radius were 0.1 N/m and 70 nm, respectively, parameters well-suited for probing soft biological samples.

Prior to each experiment, laser alignment on the cantilever tip was performed to ensure accurate deflection measurements. Probe calibration was conducted in fluid media surrounding the cells to account for changes in mechanical properties due to hydrodynamic effects. Calibration involved ramping the tip on a rigid surface (a plain 60 mm culture dish containing Milli-Q water), resulting in calibrated values of spring constant (k) = 0.06 N/m, deflection sensitivity = 122.1 nm/V, and peak force tapping amplitude sensitivity = 1117 nm/V at a 1 kHz tapping frequency. The peak force amplitude was set to 300 nm to ensure precise tip-sample interaction. Cellular targets were identified using the optical microscope integrated with the ICON head. Nanoindentation was conducted in three regions, each measuring 500 × 50 square nm^80^. The three membrane regions probed were over the nucleus, cellular periphery and midway between the two. Force-separation (F–S) curves generated from tip-sample interactions were recorded and analyzed. This method relies on cantilever deflection due to forces encountered during indentation.

Ramping parameters were optimized based on prior studies^81^ with a ramp size of 5 µm, ramp rate of 1 Hz, 2048 samples per ramp, and an applied trigger threshold of 5 nN. For each dynamic time point, a minimum of 12 cells was analyzed, with at least 10 indentations performed per cell within each 500 × 50 nm² region, yielding a minimum of 120 F–S curves. Experiments were conducted at 37°C using a temperature-controlled heated stage (Lake Shore Cryotronics, Inc., Westerville, OH). Each ramp yielded an F–S curve from which mechanical parameters such as stiffness, deformation, and adhesion were extracted. The F–S curves were initially pre-processed using Bruker’s Nanoscope Analysis v1.9 software, including application of a boxcar filter for noise reduction and baseline correction. Given the comparable dimensions of the tip and cell membrane indentation depth, a Hertzian contact model was applied to the retrace segment of the curve to extract membrane stiffness.

### α Tubulin and F-Actin Staining and Quantitative Cell Circularity Analysis

*Trp53/Pten*(−/−) murine glioblastoma cells stably expressing scramble control (SCR) or two independent shRNAs targeting ATAT1 (KD1 and KD2) were seeded onto glass-bottom chamber slides (Ibidi) at 50–70% confluency. For pharmacological experiments, cells were treated with either vehicle or GM90257 (1 μM) for 48 hours prior to fixation and analysis.

For α-tubulin immunofluorescence, cells were fixed in 4% paraformaldehyde for 10 minutes at room temperature, washed with PBS, and incubated with anti–α-tubulin antibody (Cell Signaling Technology, #2125), followed by an Alexa Fluor 594–conjugated secondary antibody (Invitrogen). Nuclei were counterstained with DAPI (Thermo Fisher Scientific; 1:1000 dilution). For F-actin staining, cells were fixed and permeabilized with 0.1% Triton X-100 in PBS and stained with rhodamine–phalloidin (Cytoskeleton Inc., PHDR1; 1:500) for 10 minutes, followed by DAPI counterstaining.

For live-cell microtubule labeling, cells were incubated with SPY650-Tubulin (Cytoskeleton Inc. # CY-SC503) according to the manufacturer’s instructions and counterstained with Hoechst 33342 (1:10,000 dilution). Live imaging was conducted using a Zeiss LSM 880 confocal microscope equipped with environmental control at 37 °C and 5% CO2, utilizing a 63 X objective.

Fixed-cell images were acquired using a Zeiss LSM 880 confocal microscope with AiryScan and 10x, 20x, or 100x objectives. Drug-treated and control samples were imaged using identical acquisition settings. Cell morphology was defined based on the α-tubulin signal and quantified using circularity analysis in Arivis 4D software, with an automated cell detection and segmentation pipeline. Total fluorescence intensities for SPY650 Tubulin and rhodamine–phalloidin were quantified using FIJI and normalized to Hoechst 33342 or DAPI, respectively. Statistical analyses were performed using GraphPad Prism.

### Stathmin-GFP Transfection and Invasion Assay

*Trp53/Pten*(−/−) murine glioblastoma cells were transfected with Lipofectamine 3000 (Invitrogen) and a plasmid encoding Stathmin-GFP (Addgene #86782), following the manufacturer’s protocol. An empty vector expressing GFP alone (pLV_TurboGFP-EV) served as a negative control. After forty-eight hours, cells were detached and subjected to transwell invasion assays. Specifically, 5x104 cells were seeded into FluoroBlok Transwell inserts, and medium containing 10% fetal bovine serum was added to the lower chamber as a chemoattractant. After overnight incubation at 37 °C, non-invaded cells were removed. Inserts were washed with phosphate-buffered saline, fixed in 4% paraformaldehyde for 20 minutes at room temperature, and washed again. Nuclei were stained with DAPI, and invaded cells were imaged using a Cytation 5 multimode imaging system. Quantification was performed by counting DAPI-positive nuclei per 20x field, and statistical analyses were conducted using GraphPad Prism software.

### MVP Protein Levels in Trp53/Pten(−/−) murine glioblastoma cells

*Trp53/Pten*(−/−) cells, scramble control (SCR) or shRNAs targeting ATAT1 (KD1 and KD2) were seeded in 6-well plates at 80% confluency. Cells were lysed in ice-cold lysis buffer containing protease and phosphatase inhibitors. Lysates were centrifuged at 17,000xg for 5 minutes at 4 °C. Protein concentrations were measured using a BCA assay (Invitrogen). Equal amounts of protein were separated by SDS–PAGE and transferred to membranes for immunoblotting. Membranes were incubated overnight at 4 °C with a primary antibody against MVP (Protein Tech #16478-1-AP, 1:1000 dilution), then with a goat anti-rabbit secondary antibody. Protein bands were detected using enhanced chemiluminescence (Invitrogen). Band intensities were quantified by densitometry in FIJI, and data were processed and analyzed statistically using GraphPad Prism.

### Analysis of Primary Cilium Length by ARL13B Staining

*Trp53/Pten*(−/−) murine glioblastoma cells or GBML1 human glioblastoma cells, stably expressing either scramble control (SCR) or shRNAs targeting mouse or human ATAT1 (KD1 and KD2), were cultured to 60% confluency. Cells were washed three times with phosphate-buffered saline (PBS), fixed in 4% paraformaldehyde for 20 minutes at room temperature, and permeabilized with 0.5% Triton X-100 in PBS for 10 minutes. Primary cilia were labeled with an anti-ARL13B antibody (Abcam, ab136648; 1:2000 dilution), followed by incubation with an Alexa Fluor 594–conjugated goat anti-mouse secondary antibody (Invitrogen). Nuclei were counterstained with DAPI. High-resolution images were obtained using a Zeiss LSM 880 confocal microscope equipped with AiryScan and a 100x objective. Cilium length was measured based on the ARL13B signal using FIJI software.

### RT-qPCR Analysis of Olig2 and c-Myc Expression

*Trp53/Pten*(−/−) cells, scramble control (SCR) or shRNAs targeting ATAT1 (KD1 and KD2) were collected for RNA isolation. Total RNA was extracted using the RNeasy Mini Kit (Invitrogen) and quantified with a Nanodrop spectrophotometer (Thermo Fisher Scientific). cDNA was synthesized utilizing 500 ng of total RNA using the iScript cDNA synthesis kit (BioRad). qPCR was conducted using SYBR Green chemistry on a QuantStudio 3 thermocycler (Invitrogen) to assess the relative mRNA expression levels of Olig2 and c-Myc, with Prlp0 as the housekeeping gene. Data were analyzed using QuantStudio software, and relative expression levels were determined using the ΔΔCt method.

### Quantification of Focal Adhesions by Vinculin Immunostaining

*Trp53/Pten*(−/−) cells, scramble control (SCR), and shRNAs targeting ATAT1 (KD1 and KD2) were seeded at 50% confluency. After fixation with 4% paraformaldehyde, cells were incubated with an anti-vinculin primary antibody (Sigma-Aldrich, V9131), followed by an Alexa Fluor 488–conjugated secondary antibody. F-actin was labeled with rhodamine–phalloidin (Cytoskeleton Inc., PHDR1; 1:500) for 10 minutes, and nuclei were counterstained with DAPI. Images were acquired using a Zeiss LSM 880 confocal microscope equipped with a 63x objective. Focal adhesion number and length were quantified from vinculin immunoreactivity using Arivis 4D software and the particle analysis pipeline.

### Analysis of DNA Content for Cell Cycle Determination

*Trp53/Pten*(−/−) cells, scramble control (SCR) or shRNA targeting ATAT1, were harvested and fixed with 4% paraformaldehyde. Following fixation, cells were permeabilized using 0.1% Triton X-100 in PBS and stained with DAPI (Invitrogen; 1:1000 dilution) to label DNA. Cell cycle profiles were obtained using an Attune flow cytometer (Invitrogen), with 50,000 events recorded per sample. Data was processed and analyzed with FlowJo software. All experiments were conducted in triplicate to ensure reproducibility and reliability.

### Data mining of The Cancer Genome Atlas (TCGA)

RNA sequencing data for high-grade glioma were obtained from TCGA *via* the cBioPortal for Cancer Genomics (http://www.cbioportal.org). The dataset included RNA-seq profiles from 261 samples, comprising 156 high-grade glioma tumor samples and 5 normal brain tissues. To investigate ATAT1 mediated target genes, we further divided the 156 samples into two groups based on their ATAT1 expression values; the ATAT1-high group contained 39 primary GBM samples constituting the tumors with the top 25% levels of ATAT1 expression; and 39 primary GBM samples constituting the tumors with the bottom 25% levels of ATAT1 expression. Raw count data from TCGA were processed and analyzed using the *edgeR* package (version 3.40.2) in R. Genes with low expression were filtered out, retaining only those with counts per million (CPM) above a defined threshold in a minimum number of samples. Normalization was performed using the trimmed mean of M-values (TMM) method to account for compositional differences between libraries. A negative binomial generalized linear model (GLM) was fitted to the count data, and differential expression between groups was assessed using likelihood ratio tests. Genes with a false discovery rate (FDR) < 0.05 were considered significantly differentially expressed.

### Statistical analysis

Data from repeated measurements are depicted as the mean ± 1 standard deviation. For quantification of Western blot intensities, at least three biological replicates were performed, and statistical significance was determined using pairwise, two tailed t tests with significance defined as p<0.05. For the Kaplan Meier survival curves in **Fig. 7**, sample size was 10 mice per group. For the cell viability measurements, experiments were performed on 96 well plates for each condition, and significance was determined with two tailed t tests as above. Fluorescence ratio measurements in were performed on at least 4 biological replicates and significance determined by two tailed t tests. Growth kinetics were determined by linear least squares fitting, and each measurement represents the mean ± 1 SD of 8 biological replicates. Statistical significance of differences in slopes of the resulting curves, corresponding to growth rates, was determined by sequential two tailed t test with significance set at p<0.05. Comparisons between dose response curves were performed with the extra sum of squares F test from Prism 10. Significance of linear regressions were performed with Spearman’s rank correlation in Prism 10.

## ACKNOWLEDGEMENTS

S.S.R. is supported by NIH grants NS073610, NS118513, NS119714, CA278293, CA210910, and CA295558 and by a Translational Adult Glioma Award from the Ben and Catherine Ivy Foundation. RSK is supported by NS118513. P.C. is supported by NS118513, CA251313, and CA250481. C.A.M. is supported by MH136567, DA049544, NS119714, AA028727, NS096833, and the William Potter Glioblastoma Research Fund. CAM and TMK are supported to UH3NS096833. T.M.K is supported by MH136567. A.Q.H. is supported by NS129671, CA282451, CA284268, a Distinguished Mayo Clinic Investigator Award, The Monica Flynn Jacoby Endowed Chair, The William J. and Charles H. Mayo Professorship, the Florida Department of Health Cancer Research Chair Fund, the Richard and Lauralee Uihlein Neuro-oncology Research Fund, the Jacquie Lorraine Goldman Fund in support of the Mayo Clinic BRIDGE Biobank, and the Florida State Funds for the Casey DeSantis Cancer Research Program. D.M. is supported by HL140411 and NS129671. We wish to thank Dr. Justin Lathia (Cleveland Clinic Lerner College of Medicine) for his gift of the L1 human GBM cell line.

## AUTHOR CONTRIBUTIONS

Conception and design: NZ, AD, SB, TK, MA, AH, NKHN, PS-M, RSK, PC, TMK, SSR Development of methodology: NZ, AD, SB, TK, MA, AH, NKHN, PS-M, RSK, PC, TMK, DM, SSR Acquisition of data: AH, NZ, AD, SB, TK, MA, AH, NKHN, PS-M, RSK, PC, TMK, SSR Analysis and interpretation of data: YL, NZ, AD, SB, TK, MA, RSK, PC, TMK, CAM, SSR

---

**Figure S1.**
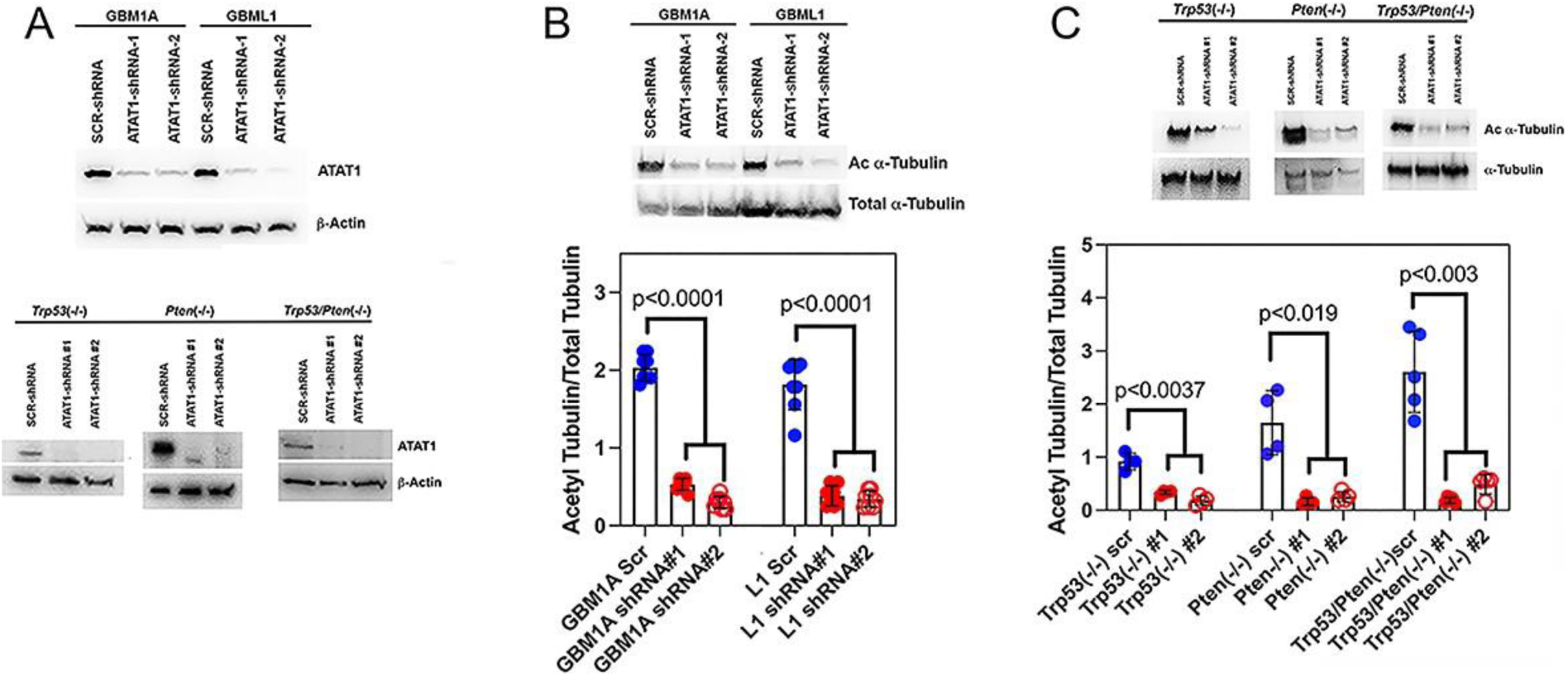
(related to Figure 1): ATAT1 knockdown reduces tubulin acetylation. (**A**): ATAT1 shRNAs reduce ATAT1 expression in both human (GBM1A and L1) and murine (*Trp53*(-/-), *Pten*(-/-) and *Trp53/Pten*(-/-)) GBM lines. (**B,C**): ATAT1 KD in human GBM1A and L1 GBM (**B**) and murine *Trp53*(-/-), *Pten*(-/-) and *Trp53/Pten*(-/-) (**C**) lines reduces acetylated tubulin levels >85%.

**Figure S2.**
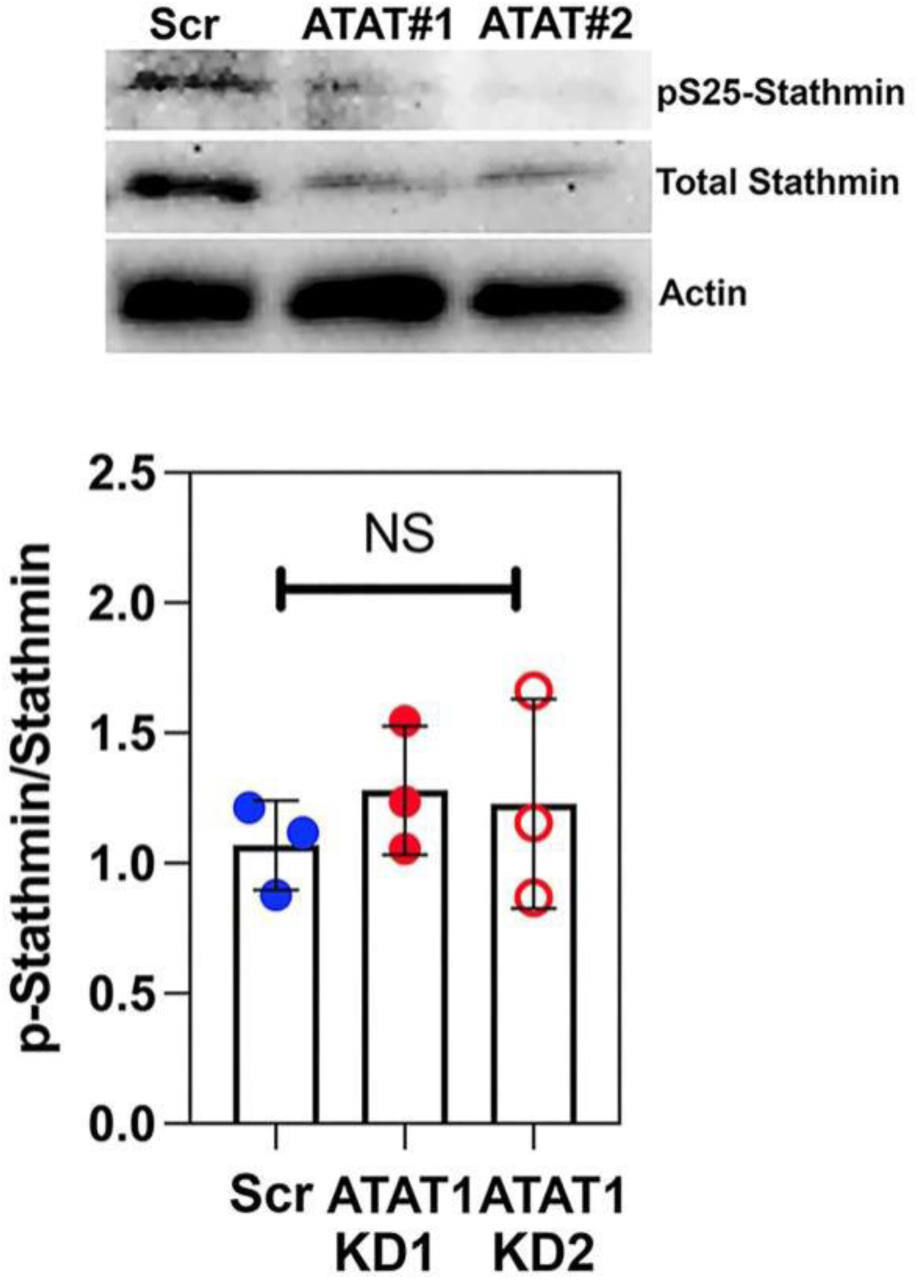
(related to Figure 1): Western blot of phosphorylated stathmin shows that ATAT1 knockdown has no significant.

**Figure S3.**
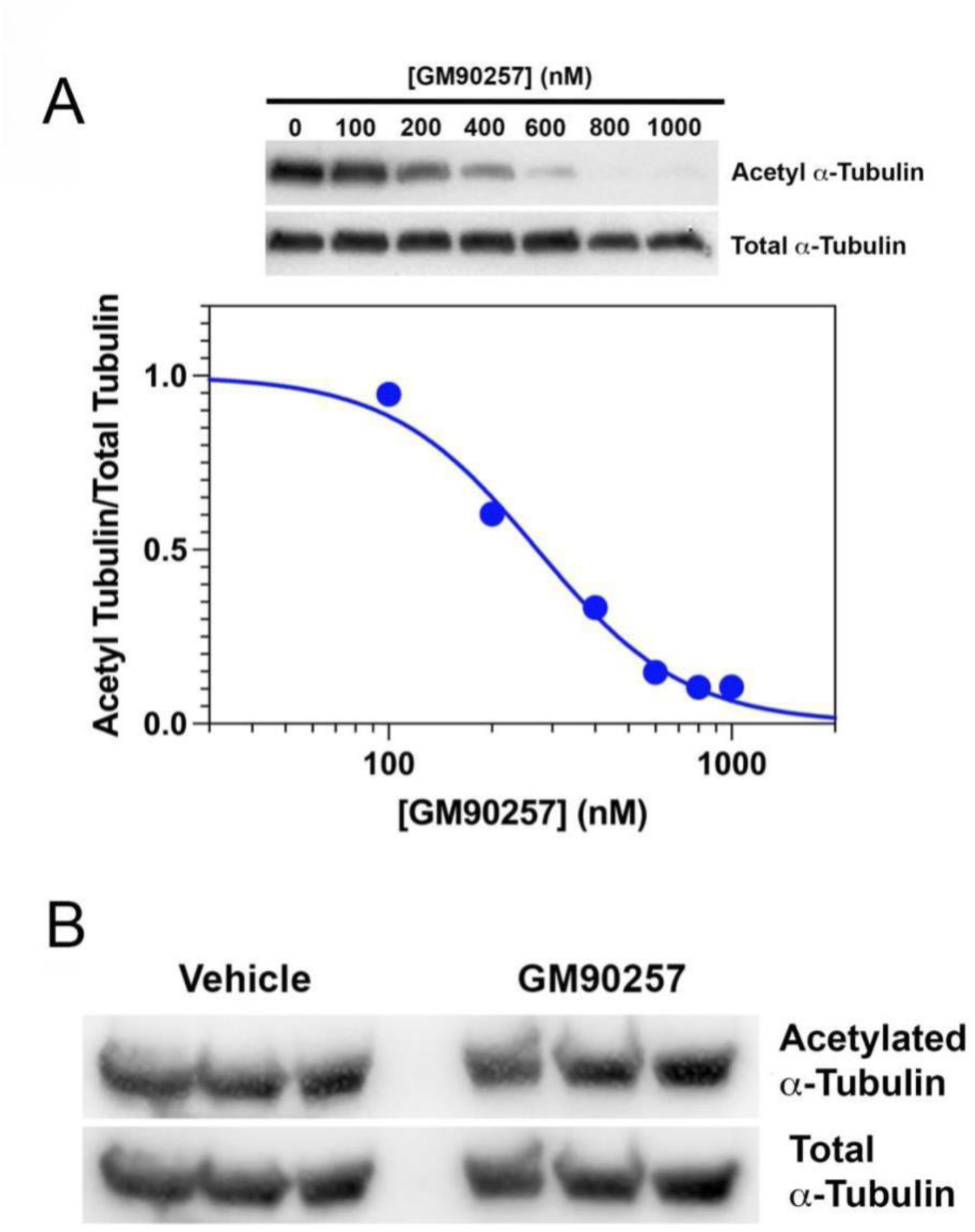
(related to Figure 1): GM90257 inhibits tubulin acetylation *in vitro*. (**A**). Dose response relationship between [GM90257] and tubulin acetylation in murine *Trp53/Pten*(-/-) GBM cells reveals an EC50 for this drug of 274 ± 69 nM. (**B**). C57Bl6 mice were injected intraperitoneally with 25 mg/kg GM90257 every two days for a total of 7 doses, were then sacrificed, and brain lysates were examined bt Western blot for levels of acetylated tubulin.

**Figure S4.**
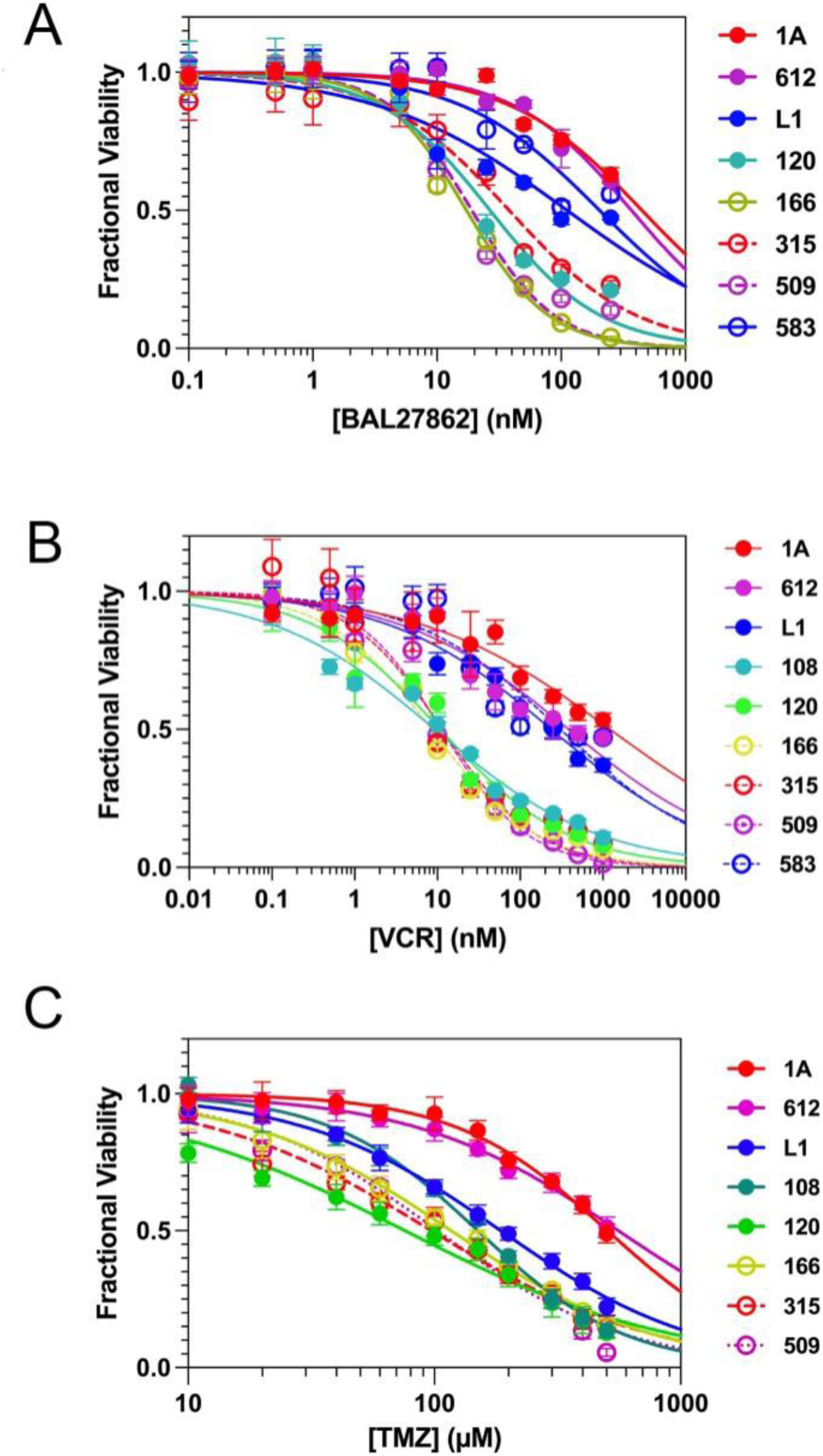
(related to Figure 2): Dose response curves for a set of primary human GBM lines. (**A-C**): Cells were treated with various concentrations of BAL27862 (**A**), Vincristine (**B**) or Temozolomide (**C**) for 72 hours, and cell count was measured with CellTiter Glo. Data are presented as cell count normalized to vehicle control versus drug concentration.

**Figure S5.**
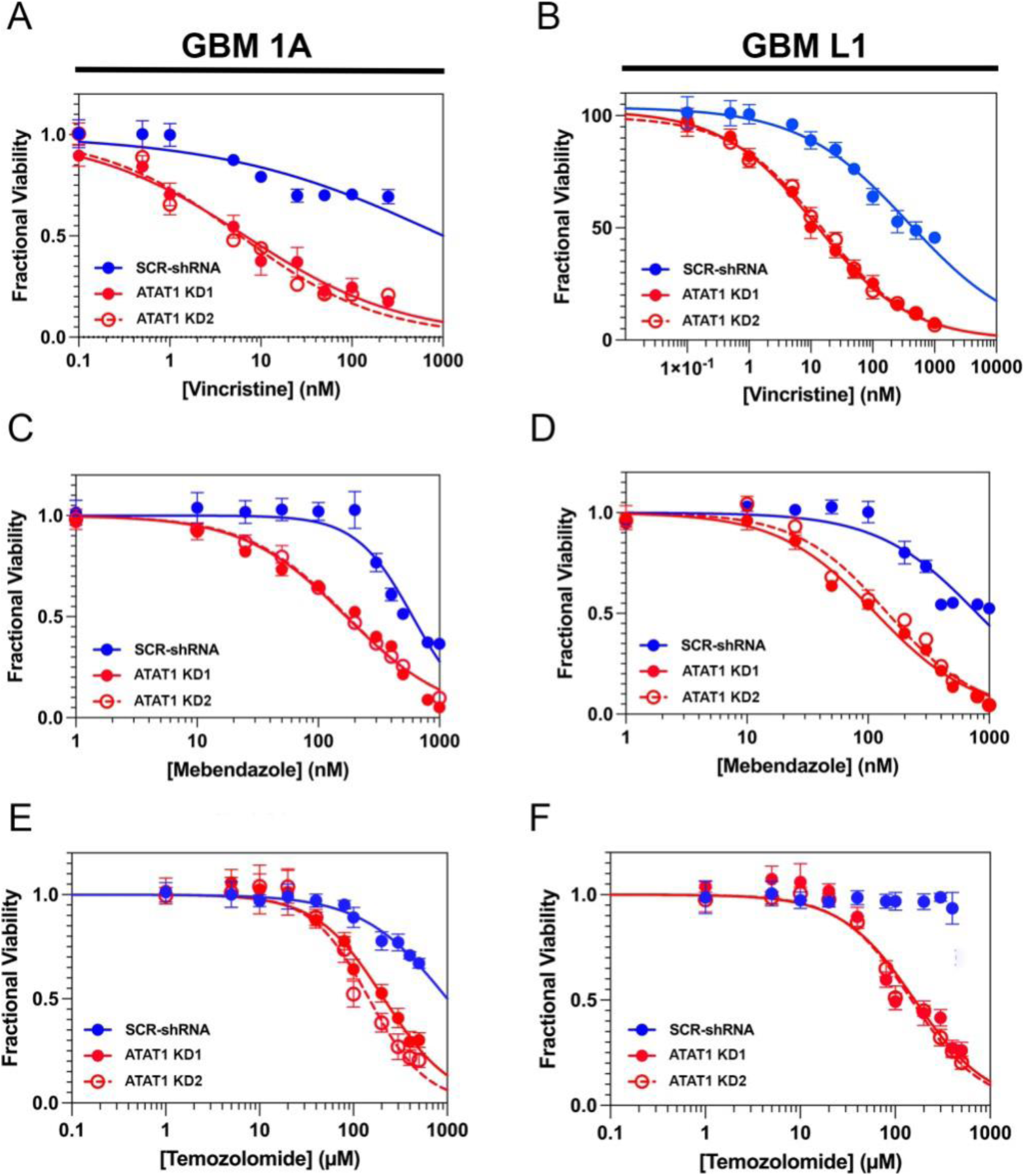
(related to Figure 2): Dose response curves for human GBM lines GBM1A (panels A,C,E) and GBML1 (panels B,D,F) in the presence (*red*) or absence (*blue*) of ATAT KD. (**A,B**): Dose response of the cell lines to vincristine. (**C,D**): dose response of the cell lines to mebendazole. (**E,F**): dose response of the cell lines to temozolomide.

**Figure S6.**
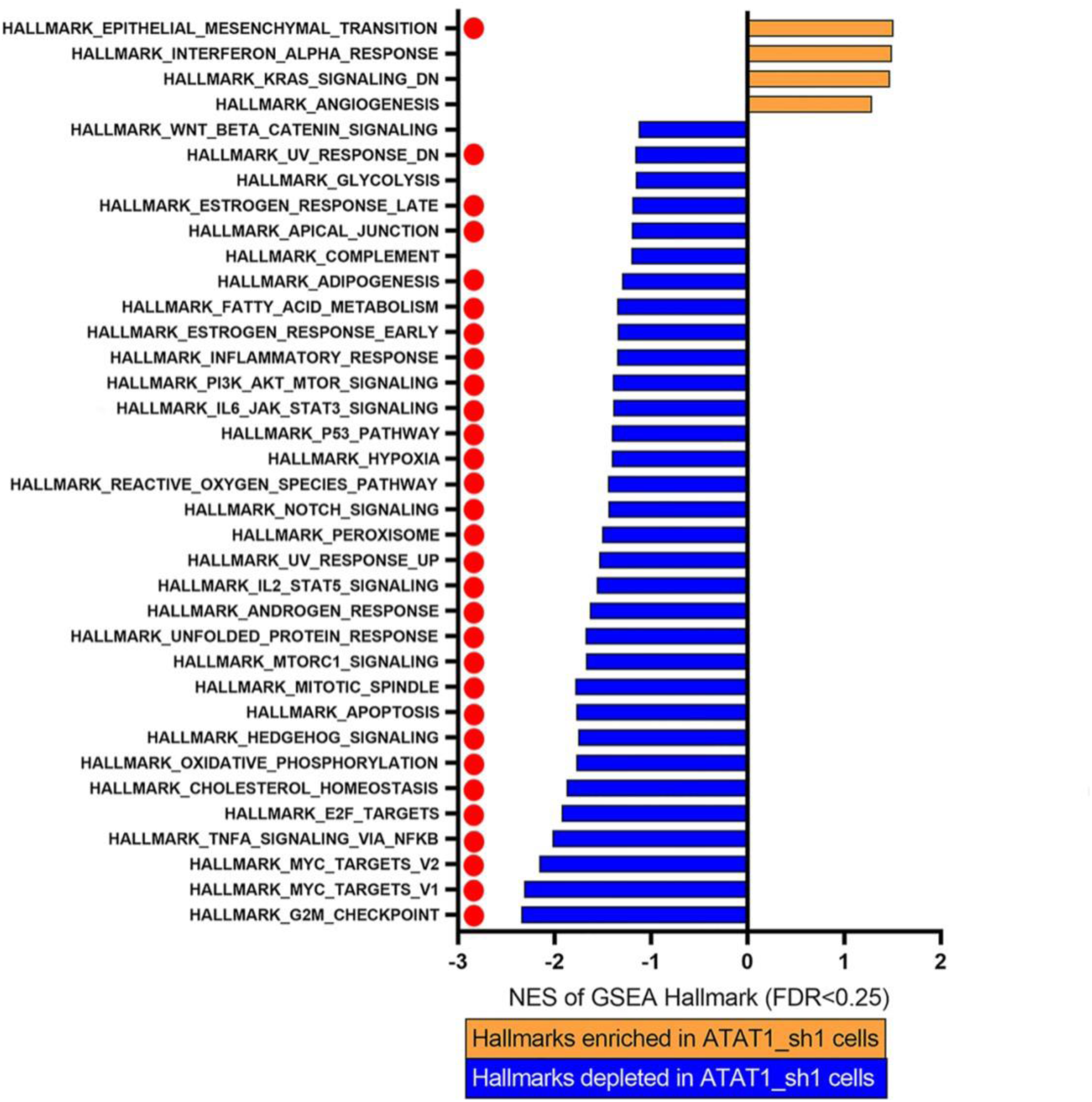
(related to Figure 3): GSEA plot for gene sets differentially regulated with ATAT1 #2 shRNA knockdown. The solid red dots indicate gene ontologies that are differentially expressed in both ATAT1 #1 (Figure 3A) and ATAT1 #2 (this figure) shRNAs knockdowns.

**Figure S7.**
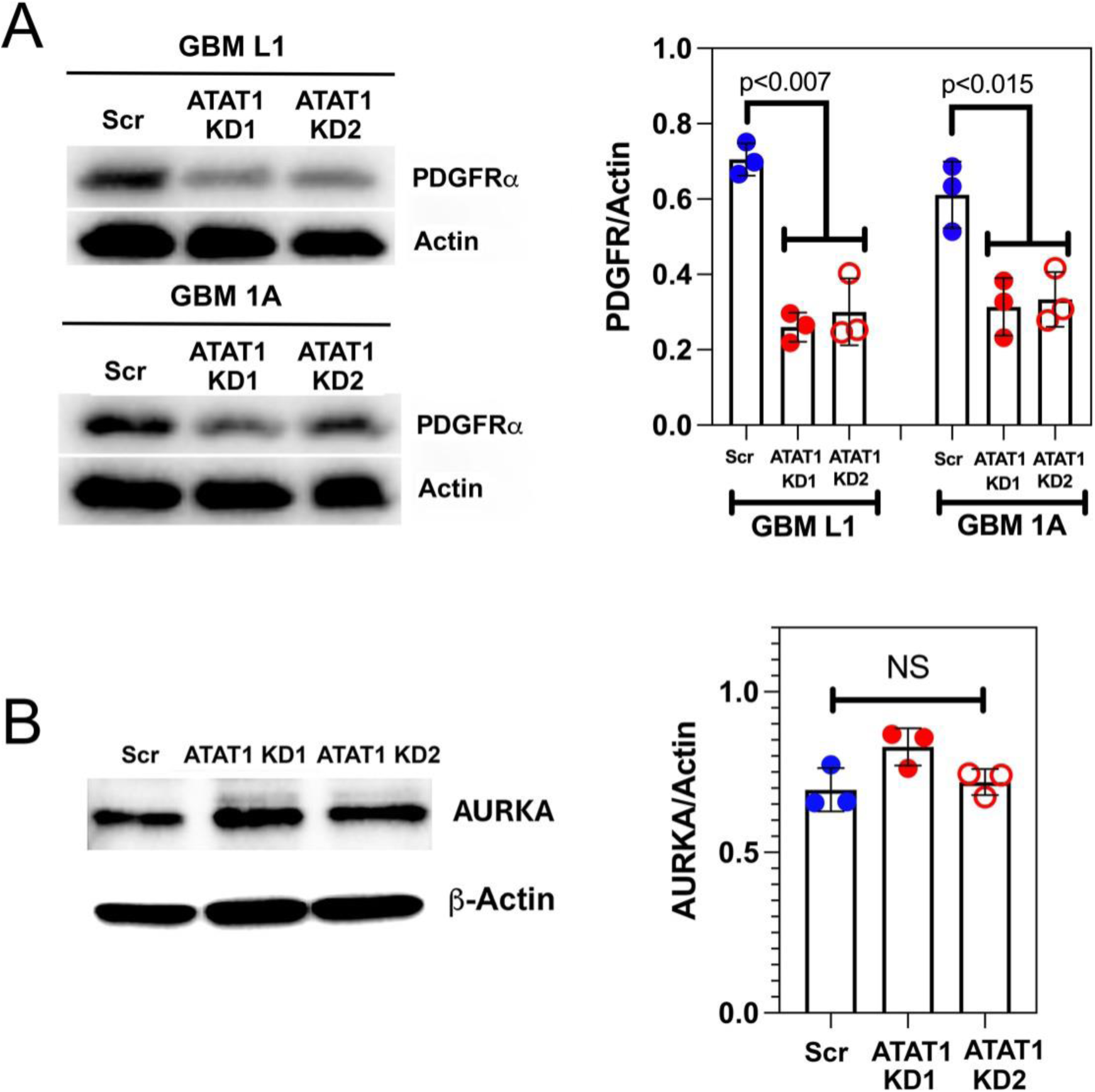
(related to Figure 4): (**A**). ATAT1 knockdown reduces expression of PDGFRa 2-3-fold in GBML1 and GBM1A. (**B**). ATAT1 knockdown in Trp53/Pten(-/-) cells has no effect on expression of Aurora Kinase A.

**Figure S8.**
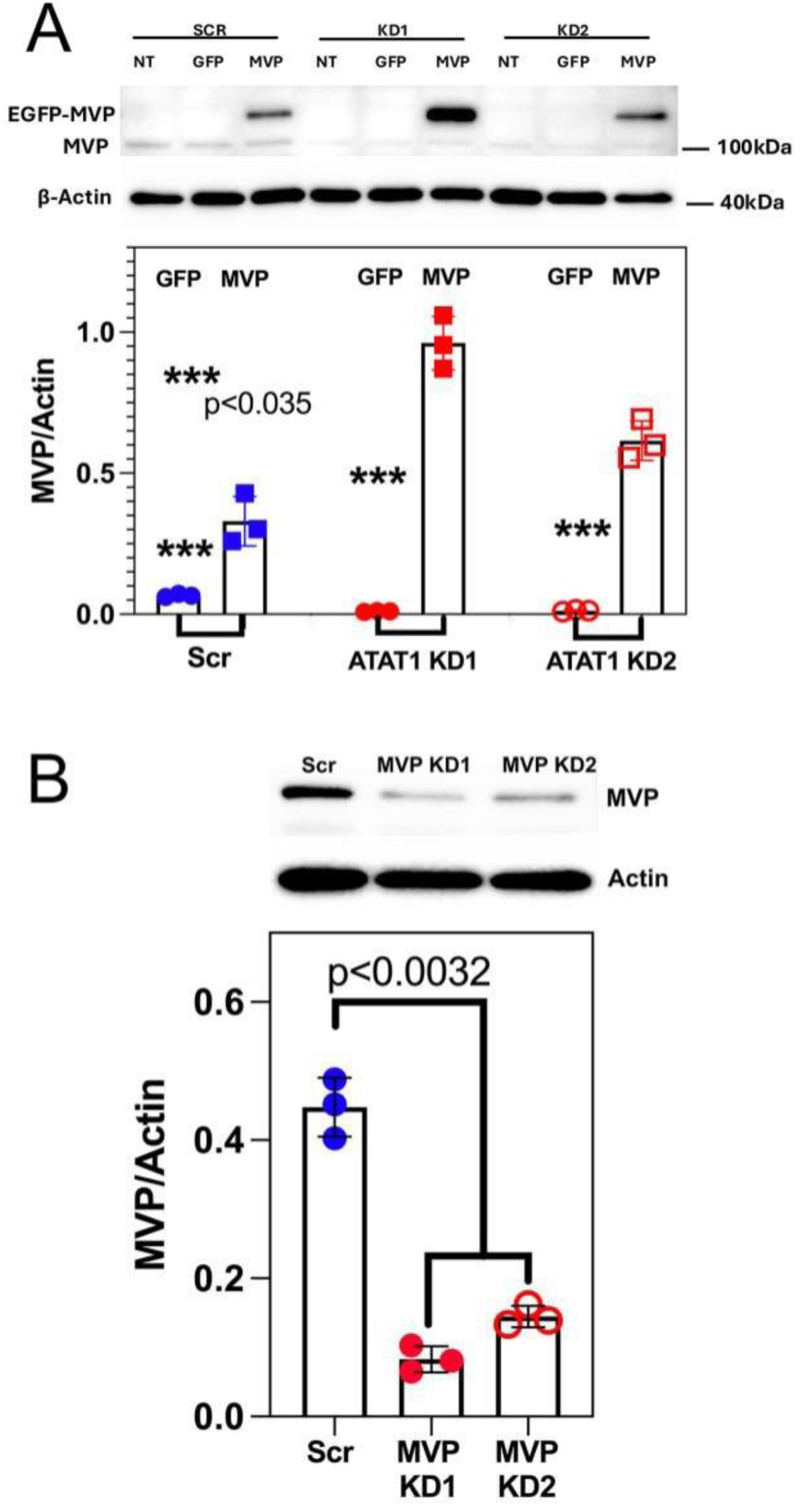
(related to Figure 6): (**A**). *Upper panel*: ATAT1 knockdown (KD1 or KD2) or Scr shRNA-treated *Trp53/Pten*(-/-) murine GBM cells were either not transfected (*NT*) or transfected with a EGFP-encoding lentiviral vector (*GFP*) or with one that encodes for an EGFP-MVP fusion protein (*MVP*). Western blots using anti-MVP antibodies demonstrate an MVP reactive band at the M_r_ expected for an EGFP-MVP fusion protein along with faint staining of endogenous MVP in cells transfected with the MVP lentivirus. *Lower panel:* quantitation of MVP immunoreactivity Scr (*blue*), ATAT1 KD1 (*closed red*), or ATAT1 KD2 (*open red*) shRNA treated cells that had been transfected with an EGFP (*circles*) or EGFP-MVP (*boxes*) encoding lentivirus. (**B**): Effect of control, Scr shRNA (*blue*) or of either of two MVP-suppressing shRNAs (*red*) in *Trp53/Pten*(-/-) murine GBM cells on MVP expression.

**Figure S9.**
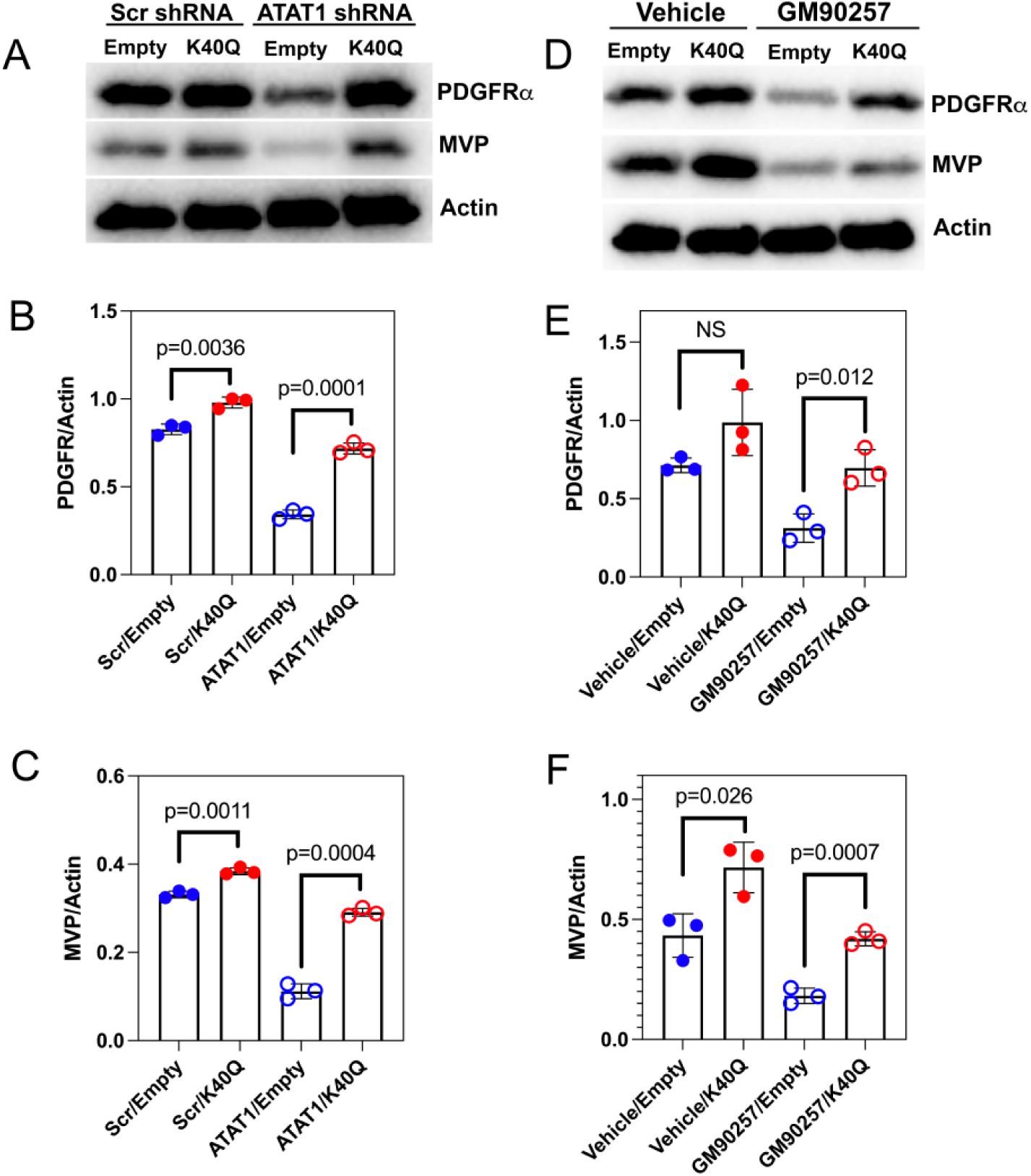
(related to Figure 6): Expression of K40Q α tubulin reverses the effects of inhibiting K40 acetylation with ATAT1 suppression (A-C) or GM90257 (D-F) on expression of PDGFRA and MVP. (**A**): Western blots of Scr and ATAT1 shRNA treated *Trp53/Pten*(-/-) cells transfected with a non-coding plasmid (“*Empty*”) or with a plasmid encoding K40Q α tubulin (“*K40Q*”). (**B**): Quantification of the Western blot intensities in (**A**) demonstrates that expression of K40Q enhances PDGFRA content by ∼10% in Scr shRNA treated cells but increases it ∼2-fold in cells suppressed for ATAT1. (**C**). Expression of K40Q enhances MVP content by ∼15% in Scr shRNA treated cells but increases it ∼3-fold in cells suppressed for ATAT1. (**D**). Western blots of *Trp53/Pten*(-/-) cells transfected with a non-coding plasmid (“*Empty*”) or with a plasmid encoding K40Q α tubulin (“*K40Q*”) and treated with vehicle or 1000 nM GM90257 for 24 hours. (**E**). Quantification of the Western blot intensities in (**D**) demonstrates that expression of K40Q enhances PDGFRA content by ∼30% in vehicle treated cells but increases it ∼3-fold in cells treated with GM90257. (**F**). Expression of K40Q enhances MVP content by 2-3 fold in both vehicle and GM90257-treated cells.

**Table S1:** Fitting Parameters of Dose Response Curves for BAL27862, Vincristine, and Temozolomide for Primary Human GBM Lines (± 1 SEM)

**BAL27862 (related to Fig. S4A)**
| *Cell Line* | *EC<sub>50</sub> (μM)* |
| --- | --- |
| 1A | 447 ± 85 |
| 612 | 368 ± 68 |
| L1 | 112 ± 24 |
| 120 | 27 ± 3 |
| 166 | 17 ± 2 |
| 315 | 38 ± 6 |
| 509 | 19 ± 2 |
| 583 | 204 ± 52 |

**Vincristine (related to Fig. S4B)**
| *Cell Line* | *EC<sub>50</sub> (μM)* |
| --- | --- |
| 1A | 1.2 ± 0.4 |
| 612 | 0.4 ± 0.08 |
| L1 | 0.2 ± 0.04 |
| 108 | 0.009 ± 0.001 |
| 120 | 0.01 ± 0.008 |
| 166 | 0.008 ± 0.001 |
| 315 | 0.01 ± 0.002 |
| 509 | 0.01 ± 0.002 |
| 583 | 0.3 ± 0.07 |

**Temozolomide (related to Fig. S4C)**
| *Cell Line* | *EC<sub>50</sub> (μM)* |
| --- | --- |
| 1A | 504 ± 79 |
| 612 | 558 ± 51 |
| L1 | 186 ± 7 |
| 108 | 150 ± 6 |
| 120 | 76 ± 6 |
| 166 | 117 ± 6 |
| 315 | 97 ± 7 |
| 509 | 103 ± 7 |

**Table S2:**
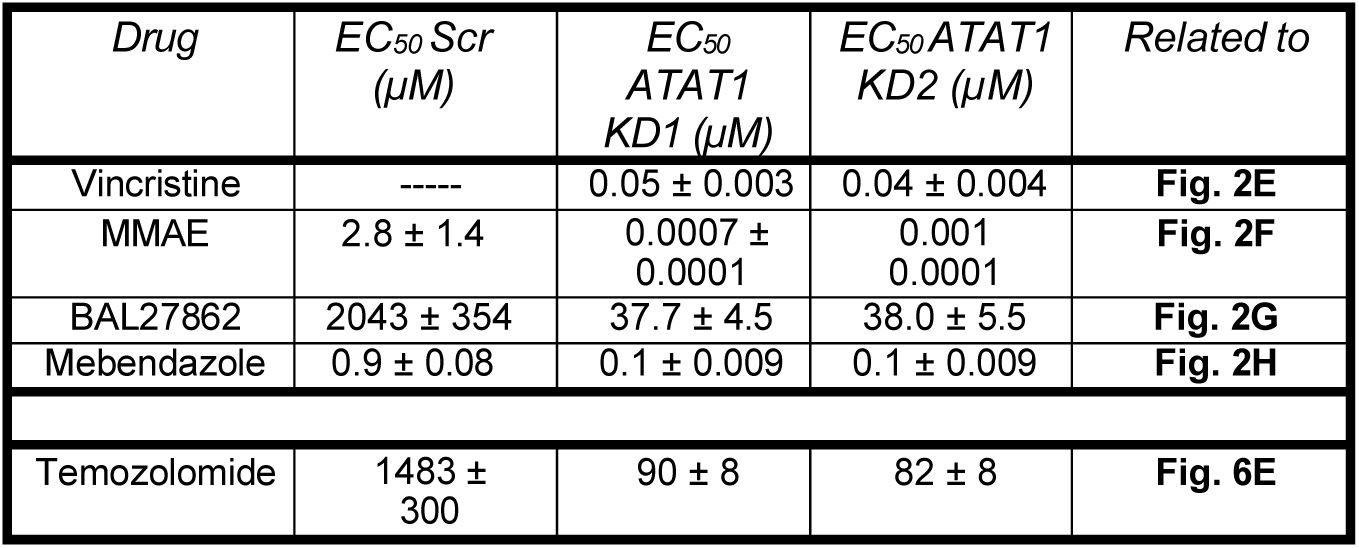
Effect of ATAT1 KD on EC_50_ of Microtubule Depolymerizers and Temozolomide in Murine *Trp53/Pten*(-/-) Cells (± 1 SEM)

**Table S4:**
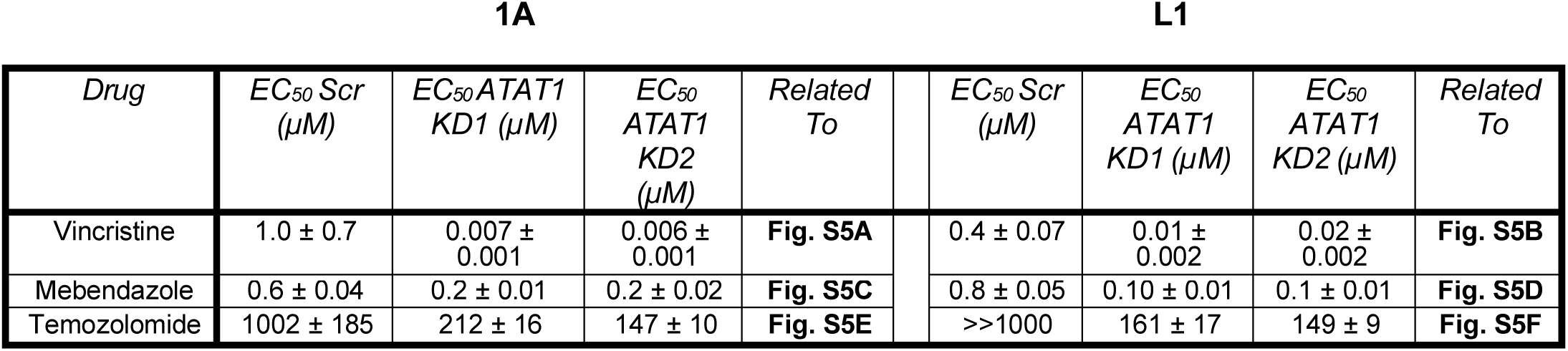
Effect of ATAT1 KD on EC_50_ of Microtubule Depolymerizers and of Temozolomide in Human L1 and 1A GBM Cells (± 1 SEM)

**Table S5:** Effect of MVP Transfection or Knock Down on Temozolomide and Mebendazole Potency in Murine *Trp53/Pten*(-/-) and Human 315 Cells (± 1 SEM)

| *Treated With:* | *Cell Line* | *Drug* | *EC<sub>50</sub> (μM)* | *Related To* |
| --- | --- | --- | --- | --- |
| Scr + Empty Vector | *Trp53/Pten*(-/-) | Temozolomide | 835 ± 108 | **Fig. 6D** |
| ATAT1 KD1+ Empty Vector | *Trp53/Pten*(-/-) | Temozolomide | 85 ± 55 | **Fig. 6D** |
| ATAT1 KD2+ Empty Vector | *Trp53/Pten*(-/-) | Temozolomide | 88 ± 8 | **Fig. 6D** |
| Scr + MVP | *Trp53/Pten*(-/-) | Temozolomide | 1405 ± 248 | **Fig. 6D** |
| ATAT1 KD1 + MVP | *Trp53/Pten*(-/-) | Temozolomide | 371 ± 40 | **Fig. 6D** |
| ATAT1 KD2 + MVP | *Trp53/Pten*(-/-) | Temozolomide | 449 ± 42 | **Fig. 6D** |
| Scr | *Trp53/Pten*(-/-) | Temozolomide | 1030 ± 342 | **Fig. 6E** |
| MVP KD1 | *Trp53/Pten*(-/-) | Temozolomide | 80 ± 8 | **Fig. 6E** |
| MVP KD2 | *Trp53/Pten*(-/-) | Temozolomide | 97 ± 5 | **Fig. 6E** |
| Scr | 315 | Temozolomide | 104 ± 5 | **Fig. 6F** |
| MVP KD1 | 315 | Temozolomide | 25 ± 2 | **Fig. 6F** |
| MVP KD2 | 315 | Temozolomide | 23 ± 2 | **Fig. 6F** |
| MVP KD3 | 315 | Temozolomide | 21 ± 2 | **Fig. 6F** |
| Scr | *Trp53/Pten*(-/-) | Mebendazole | 468 ± 29 | **Fig. 6G** |
| MVP KD1 | *Trp53/Pten*(-/-) | Mebendazole | 60 ± 4 | **Fig. 6G** |
| MVP KD2 | *Trp53/Pten*(-/-) | Mebendazole | 89 ± 1 | **Fig. 6G** |

